# Template Learning: Deep Learning with Domain Randomization for Particle Picking in Cryo-Electron Tomography

**DOI:** 10.1101/2024.03.20.585905

**Authors:** Mohamad Harastani, Gurudatt Patra, Charles Kervrann, Mikhail Eltsov

**Affiliations:** Department of Integrated Structural Biology, Institute of Genetics and Molecular and Cellular Biology, Illkirch, France; Inria Center at University of Rennes, SAIRPICO Team, Cellular and Chemical Biology Unit, U1143 INSERM, UMR3666 CNRS, Institut Curie, PSL Research University, Campus universitaire de Beaulieu, Rennes Cedex, France

**Keywords:** Cryo-ET, Particle Picking, Deep Learning, Domain Randomization, Physics-based Simulations, Cell Crowding, Molecular Mechanics

## Abstract

Cryo-electron tomography (cryo-ET) enables the three-dimensional visualization of biomolecules and cellular components in their near-native state. Particle picking, a crucial step in cryo-ET data analysis, is traditionally performed by template matching—a method utilizing cross-correlations with available biomolecular templates. Despite the effectiveness of recent deep learning-based particle picking approaches, their dependence on initial data annotation datasets for supervised training remains a significant limitation. Here, we propose a technique that combines the accuracy of deep learning particle identification with the convenience of the model training on biomolecular templates enabled through a tailored domain randomization approach. Our technique, named Template Learning, automates the simulation of training datasets, incorporating considerations for molecular crowding, structural variabilities, and data acquisition variations. This reduces or even eliminates the dependence of supervised deep learning on annotated experimental datasets. We demonstrate that models trained on simulated datasets, optionally fine-tuned on experimental datasets, outperform those exclusively trained on experimental datasets. Also, we illustrate that Template Learning used as an alternative to template matching, can offer higher precision and better orientational isotropy, especially for picking small non-spherical particles. Template Learning software is open-source, Python-based, and GPU and CPU parallelized.

## Introduction

Recent technological advances in cryo-electron tomography (cryo-ET) expanded our capabilities to explore the three-dimensional (3D) cell architecture at the molecular scale. On one side, the automation of vitreous cryo-lamellae milling significantly enhanced the throughput of tomographic sample preparation ^1–3^. On the other side, improvements in the hardware of transmission electron microscopes (TEM), especially new generations of direct electron detectors, enabled the routine acquisition of tomograms with the resolution that allows identification of biomolecules and determination of their structure and its variations, directly in their functional environment ^4–8^.

The cryo-ET pipeline begins with the vitrification of the sample by plunge or high-pressure freezing. In the case of bulky samples such as eukaryotic cells and tissues, this step is followed by thinning using cryo-focused ion beam milling or cryo-sectioning. A series of 2D images (micrographs) is then obtained by rotating the sample in cryo-TEM. The resulting images, referred to as a tilt series, are computationally reconstructed into a 3D volume called a tomogram ^9,10^.

Although cryo-ET allows the acquisition of high-resolution data from cellular samples, it faces serious limitations. Existing sample preparation and cryo-TEM setups limit the tilting range, typically within ±60 degrees. This angular limitation leads to a missing wedge region in Fourier space resulting in severe real-space data anisotropies commonly referred to as missing wedge artifacts ^11–14^. In addition, the sensitivity of biological samples to radiation damage requires image acquisition with a limited electron dose, resulting in a compromised signal-to-noise ratio (SNR).

Locating copies of biomolecules in cryo-electron tomograms, known as particle picking, is important for understanding biomolecular distribution in cells and constitutes an initial stage in obtaining higher resolution 3D structures via averaging and classification ^15,16^. However, manual particle picking is labor-intensive, further complicated by the challenges posed by the missing wedge and low SNR.

A well-established method to automate particle picking using the similarity with an external template via cross-correlation is called template matching ^17,18^. In recent years, several deep learning-based methods have emerged and were shown capable of outperforming template matching, for some studies, in speed and accuracy ^19–24^.

A specialized track of the SHape REtrieval Contest (SHREC) was launched to compare various deep learning methods for cryo-ET particle picking ^25^. These evaluations were conducted on simulated datasets, where tomograms with ground truth annotations are available and are used for training. However, on experimental datasets, supervised deep learning methods will require a training dataset, usually obtained via manual template matching-assisted picking.

Open databases for 3D structural data of biomolecules, e.g., the Protein Data Bank (PDB) ^26^, give access to structures that can serve as templates for template matching. However, despite the remarkable advancements in the physical modeling of cryo-ET data ^27–30^, templates were not used for supervised training of deep learning models on customized particle picking ^20,25,31^. Some methods found a use for simulated datasets in training deep learning-based general-purpose particle picking (e.g. cryoYOLO ^32^) or unsupervised structural mining (e.g., TomoTwin ^31^). However, these methods face several limitations, including molecular crowding, structural variations, or computational complexity ^22,23,31,33^.

Using simulations derived from previously defined or inferred structures, such as those obtained from *in vitro* or *in silico* studies, to train models to annotate similar structures in cryo-electron tomograms would be highly advantageous over the time-consuming initial annotation directly from *in situ* data. This raises the question of why such methodologies are not more widespread. The literature points to a challenge known as the “synthetic-to-real domain gap” ^34^ where deep learning models trained on simulations can only operate in the synthetic domain. This domain is characterized by attributes like texture, illumination, noise, background, and other factors, which are notably different from those in the real-world domain ^34^.

Domain randomization is a state-of-the-art approach for addressing the domain gap during synthetic dataset generation, specifically in the context of training models for object classification and segmentation. Domain randomization is based on simulating non-realistic scenarios that were proven powerful to supervised deep learning of models capable of generalizing to real-world data ^35,36^. It hinges on three principles: 1) utilizing models of target objects with randomized shape and pose variations; 2) incorporating a mix of diverse objects termed “distractors”, positioned in the background near the target objects; and 3) integrating a broad spectrum of random rendering options.

In cryo-ET literature, addressing the synthetic-to-real domain gap for deep learning training has been largely overlooked. Nevertheless, a method called CryoShift ^37^ has utilized domain adaptation and randomization techniques for subtomogram classification, primarily focusing on mapping the synthetic data domain to resemble the real data during classifier training. Yet, this method was limited to handling pre-isolated subtomograms of molecules of interest and was not designed to process entire tomograms for particle pickling or segmentation.

Here, we introduce “Template Learning”, a domain randomization-based strategy for generating cryo-ET simulated tomograms, incorporating considerations for molecular crowding, structural variabilities, and data acquisition variations, with which we can learn deep learning models to achieve state-of-the-art performance in particle picking within cryo-ET experimental tomograms. In this paper, we detail our approach and systematically benchmark it against alternative methods, utilizing a recently published, exhaustively annotated *in situ* cryo-ET dataset. We also apply Template Learning to nucleosome picking within a densely populated cryo-ET sample of isolated mitotic chromosomes. We illustrate that models trained on simulated datasets, optionally fine-tuned on experimental datasets, outperform those exclusively trained on experimental datasets. Also, we illustrate that Template Learning used as an alternative to template matching, can offer higher precision and better orientational isotropy, especially for picking small non-spherical particles.

## Results

### Customized simulations to train deep learning models on particle picking

Template Learning is a streamlined pipeline for simulating cryo-ET synthetic data for training deep learning models on particle picking. The fundamental goal of Template Learning is to train deep learning models in three aspects:

1. Identifying targeted particles across diverse variations.
2. Differentiating target particles from other structures, especially in crowded environments.
3. Expanding the capabilities of the models to handle the inherent variations present in cryo-ET experimental data.

The practical application of these aspects is achieved through our proposed pipeline, illustrated in Fig.1 and further elaborated in the subsequent sections.

**Fig. 1:**
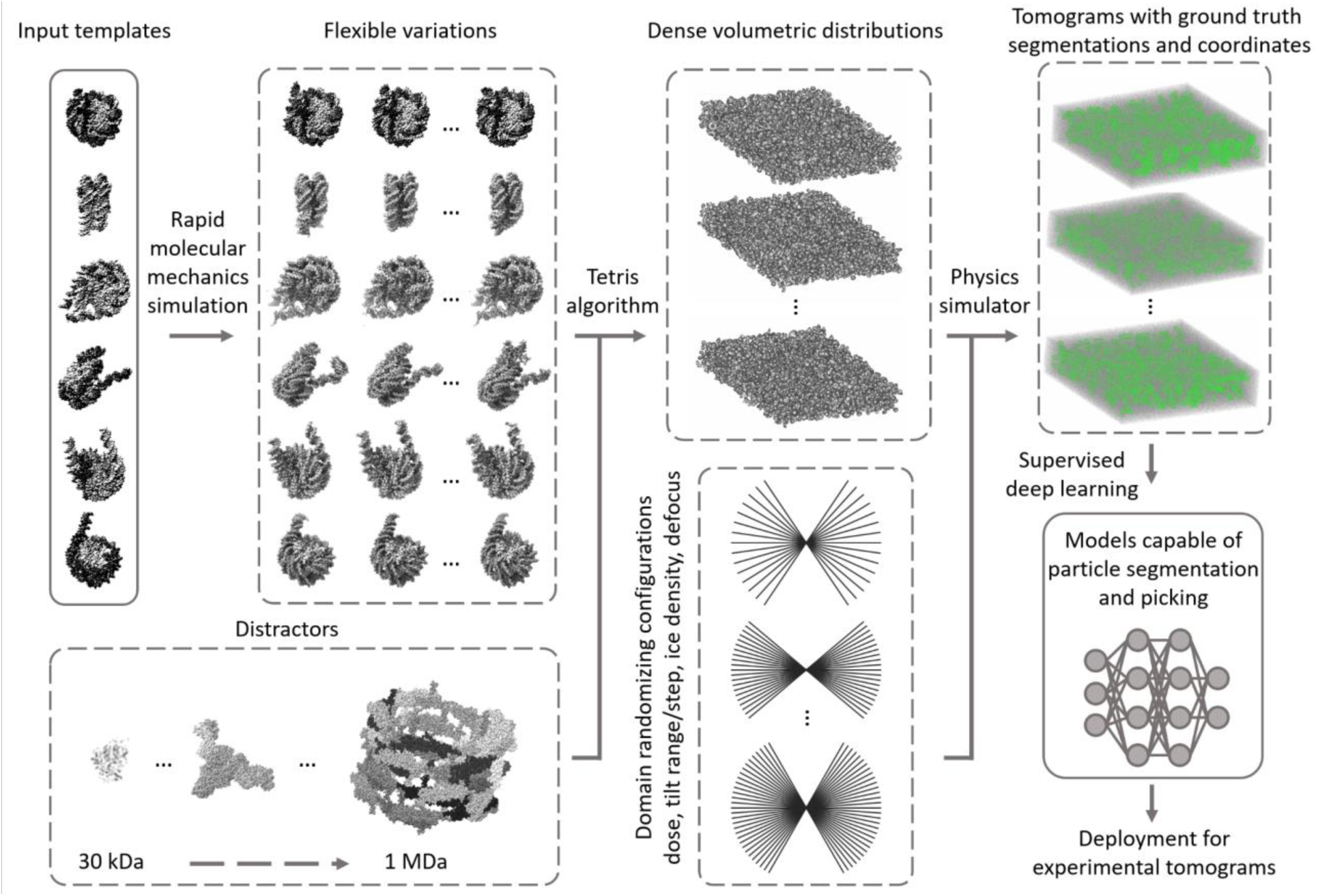
Template Learning workflow for generating simulations used for supervised deep learning used for cryo-ET picking of target particles. Template structures are used as input, commonly differing by some structural variations of the target biomolecule (we mean by the target biomolecule, the structure that the deep learning model is desired to learn to annotate in cryo-ET tomograms, in the figure it is the nucleosome). The input templates are augmented with flexible variations based on fast molecular mechanics simulations (see text for details). A general set of other proteins, different from the input templates, are used as “distractors” when generating synthetic data. The templates, along with their flexible variations, and distractors are placed at random orientations in close proximity using the “Tetris algorithm”— an algorithm proposed in this work to enable fast simulation of high molecular crowding. A physics-based simulator (Parakeet ^28^) is used to generate synthetic cryo-ET data using a set of different parameters that were chosen to produce domain randomization at the lowest cost, which are the electron dose, tilt range, tilt step, ice density, and defocus. The resulting output comprises tomograms and ground truth segmentations and coordinates corresponding to the templates. The generated data is subsequently utilized to train deep learning models (e.g., DeepFinder ^19^) for experimental tomogram segmentation and particle picking. The PDB IDs used for templates in this figure are 2PYO, 7KBE, 7PEX, 7PEY, 7XZY, and 7Y00, and for distractors are 3QM1, 7NIU, and 6UP6. The display of structures was performed using ChimeraX ^52^ and IMOD^47^ software

### Accounting for the structural and orientational variabilities

To enable trained deep learning models to effectively identify various variations of the target biomolecule, we utilize two following strategies:

1. We use multiple templates of one target biomolecule to account for compositional or significant conformational variabilities of the target particles.
2. We generate random flexible variations of each template using Normal Mode Analysis (NMA—a method for fast molecular mechanics simulations, see Methods for details) ^38^. The templates with their flexible variations are then simulated at uniformly random orientations to account for orientational variations.

Templates in this study are typically atomic structures. However, since the availability of atomic structures may vary for different target biomolecules, using lower resolution cryo-EM density maps (volumes) is also possible and is explored in a subsequent section.

### Accounting for the crowded environment

The existing literature on domain randomization for object recognition in natural images simulates the target objects within a background containing a library of other random objects ^39^. These additional objects, distractors, obscure or cast shadows on segments of the target objects. In this study, we selected 100 dissimilar protein assemblies, with diverse molecular weights ranging from 30 kDa to 1 MDa to serve as elements within the background (a comprehensive list of these proteins is available in Extended Data Fig.1). To achieve a similar effect of distractors for cryo-ET data, i.e., overlapping at the level of the tilt series and the missing wedge artifacts, these proteins must be positioned in close proximity to the templates during data modeling.

Recent works ^27,40^ that involve simulating cryo-ET data for practical applications (e.g., deep learning) utilize an iterative brute-force random placement algorithm, involving the rotation of a duplicate of a molecule in each iteration to find a suitable non-overlapping position within the sample. Nonetheless, random placement leads to unstructured empty spaces between the molecules that prevent achieving highly dense samples.

Two other approaches simulate molecular crowding of cryo-ET data ^41,42^. The first represents molecules as spheres, based on calculating the minimum bounding sphere for each molecule, and simulates crowding by optimizing a sphere-packing problem. However, this approach is limited to generating high crowding only for spherical-shaped molecules. The second builds on the first by including an additional step of molecular dynamics simulations to enhance packing density. However, routine application of this approach requires significant computational resources.

To achieve high molecular crowding while maintaining generality and efficiency, we drew inspiration for a simplified approach from a recent concept on packing generic 3D objects ^43^, a solution we term the “Tetris algorithm.” Our algorithm, illustrated in Fig. 2, operates on an intuitive principle: it places molecules iteratively, with each iteration positioning a new molecule at a uniform random orientation as close as possible to those already placed (the user only needs to choose the minimum distance). For more details on the principles of the Tetris algorithm, see Methods.

**Fig. 2:**
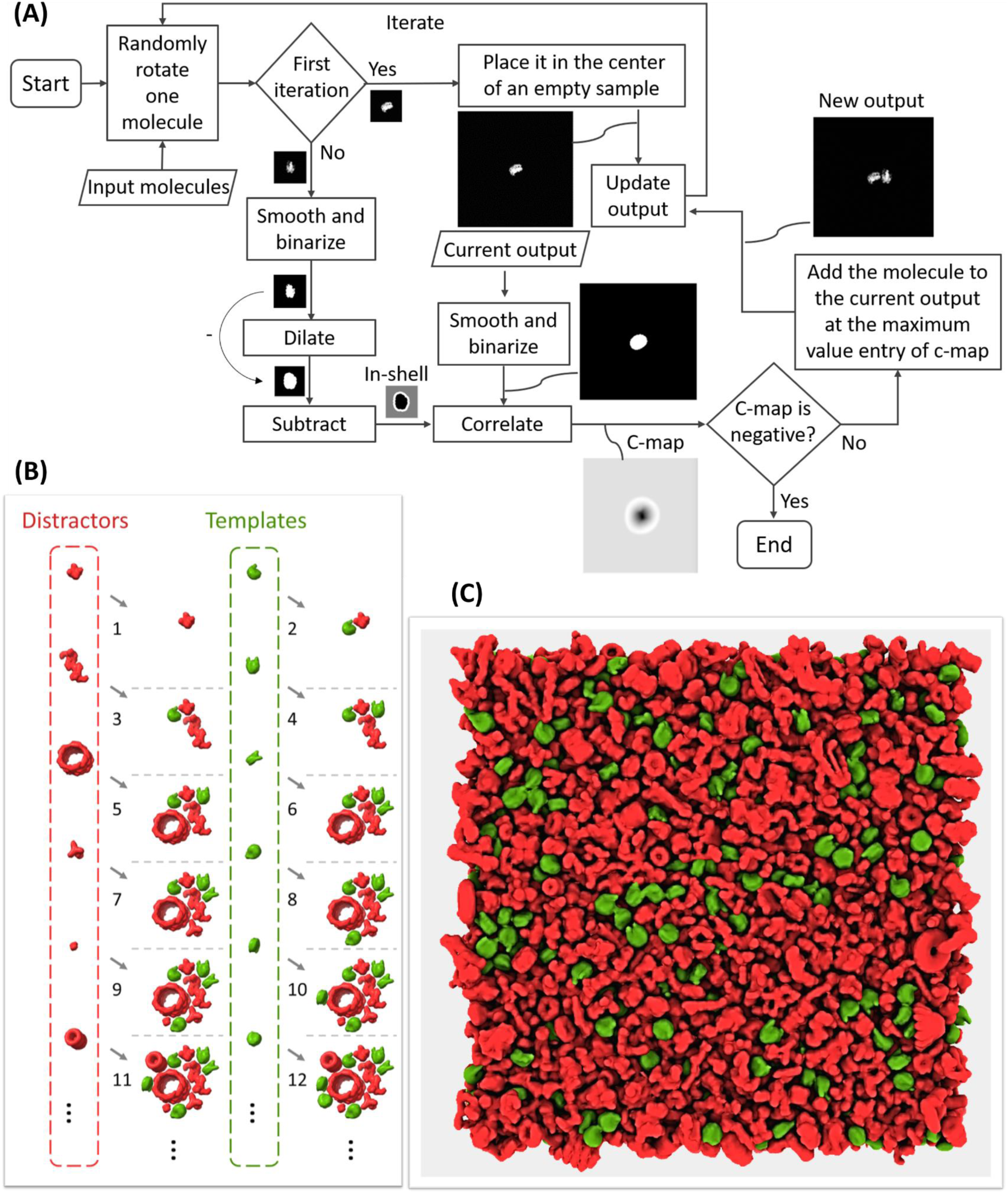
The Tetris algorithm for generating crowded 3D samples. **A** The flowchart of the algorithm, demonstrated in 2D for simplicity. The molecules that are desired to populate the sample are used as input. In the initial iteration, a molecule is positioned at the center of an expanded volumetric sample, forming the current output. In subsequent iterations, a new molecule undergoes binarization and dilation. By multiplying the binarized version with a large positive value and subtracting the result from the dilated version, an insertion shell is generated (referred to as ‘in-shell’). This insertion shell is then correlated with a binarized version of the current output volume, resulting in a correlation map (referred to as ‘c-map’). Positive values in the c-map (white voxels) represent viable positions for adding the new molecule without intersection, whereas negative and zero values (black and gray voxels, respectively) represent positions where intersection with existing molecules occurs or where the distance from other molecules is too large. If the correlation map is not entirely negative, the molecule is added to the current output at the index of the maximum entry, ensuring it avoids intersections while maintaining the highest compactness. **B** An example of the Tetris algorithm input and output at each iteration, alternating between placing templates (in green) and distractors (in red). The numbers in **B** correspond to the output at different iterations of the algorithm in **A**. **C** An example of a Tetris algorithm output in 3D.

### Accounting for the real-world domain variance

Simulations can partially replicate the domain of real-world data; however, variations in the real world can be unpredictable, and simulations simplify some complex phenomena for practical reasons. Thanks to domain randomization ^34,36^, simulating an exact match of the real world is not necessary for training deep learning models capable of generalizing to real-world data. Instead, domain randomization focuses on training these models on diverse simulated scenarios to minimize the influence of the domain on their capabilities.

In this work, we employ a cryo-ET physics-based simulator called Parakeet ^28^. Parakeet, and its MULTEM backend ^44^, provide control over numerous parameters for simulating cryo-ET hardware and the behavior of biological samples during data recording. While many parameters related to the electron beam, lenses, detector, sample, and data acquisition strategy can be controlled, it might not be necessary to randomize every available parameter. On one hand, the combinations required to cover all conditions grow exponentially with the number of variables. On the other hand, some variables might have undesirable effects on the overall simulation. Here, we propose varying a few essential parameters that significantly influence the output simulation: the electron dose, defocus, tilting range, tilting step, and ice density, while giving other parameters default values. The electron dose is crucial for varying the SNR and the irradiation effects on the sample. The defocus plays a key role in controlling the Contrast Transfer Function (CTF) ^45^. Varying the tilting range and step may not replicate experimental data acquisition strategies, however, it can expose the deep learning model to variations that increase its robustness, for example, to artifacts and imperfections resulting from tilt series alignment and reconstruction. Lastly, different ice densities can simulate the variability of sample thickness and solvent composition. A combination of three values for the defocus and two values for each of the other variables results in 48 combinations that we found sufficient for efficient domain randomization.

### Benchmarking Template Learning for picking ribosomes *in situ*

In a recent study ^20^, the first exhaustively annotated *in situ* cryo-ET dataset was published (EMPIAR-10988). Here, we use this dataset to evaluate the proposed Template Learning thoroughly.

In the following, we generate, based on the Template Learning workflow, a simulated dataset to train DeepFinder ^19^ for ribosome annotation. We compare the performance of this model (i.e., DeepFinder trained solely on simulations) to the contemporary techniques, namely, DeePiCt ^20^ and DeepFinder (trained solely on annotated experimental data), and template matching.

We organized our experiments based on two principles. First, we benchmark the complete method that integrates the aforementioned concepts of Template Learning. Then, we conducted experiments where specific elements from the data simulation pipeline were intentionally omitted (e.g., the structural variations of the target template, the crowding, etc.) to assess their impact on performance and to offer guidelines for future users. Second, acknowledging that the size of a simulated dataset is user-defined (generally in deep learning, the more training data, and the more diversity, the better), we ensured the feasibility of routine use of the method by limiting its time requirement to a maximum of two days on a single GPU (within one day on a typical computing node with 4 GPUs, see Software requirements and availability section for details).

### The typical Template Learning workflow

To establish a Template Learning workflow, we proceeded with the following steps. We selected 6 eukaryotic ribosome PDB structures (PDB IDs: 4UG0, 4V6X, 6Y0G, 6Y2L, 6Z6M, and 7CPU). The 6 selected structures differ slightly in composition or conformation. By applying NMA, we generated 25 flexible variations for each structure (refer to the NMA section in Methods for details). While generating synthetic data, we employed the aforementioned general distractors (a full list of distractors is given in Extended Data Fig.1). To simulate a crowded environment, we generated volumes from the template structures (i.e., the 6 ribosome PDBs and their flexible variations) and distractors with a sampling rate of 16 Å using Eman2 ^46^ software e2pdb2mrc. We used the generated volumes as input to populate 48 crowded volumetric distributions of size 192 x 192 x 64 voxel^3^ using the Tetris algorithm, with ribosome templates appearing at a frequency of one for every 5 distractors to produce a balanced volumetric density ratio balance between distractors and templates in the simulated tomograms (see Tetris algorithm section in Methods for details). Subsequently, we fed the sample information (i.e., atomic structures of templates and distractors with the positions and orientations of crowding generated using the Tetris algorithm) to Parakeet ^28^ to simulate 48 tilt-series using a dose symmetric tilting scheme using the parameters listed in Extended Data Tab.1. All other parameters adhered to default values for Volta Phase Plates (VPP) simulations within the Parakeet software. Subsequently, we binned the simulated tilt series to a similar sampling rate as the analyzed dataset (13 Å/pixel) and reconstructed them in IMOD ^47^ using Weighted Back Projection. We used the tomograms and their corresponding ground-truth template coordinates (around 6,500 simulated ribosomes) and segmentations (i.e., volumes of the same size as the tomograms where template instances appear in white over black background, illustrated in Extended Data Fig.2) to train DeepFinder ^19^ using its default parameters settings (the list of parameters is given in Extended Data Tab.2).

We applied the DeepFinder simulations-trained model to the 10 VPP tomograms of EMPIAR-10988 dataset without any preprocessing, resulting in segmentation maps, i.e., ribosome segmentations based on the decisions of the model. Subsequently, we employed the MeanShift algorithm offered by the DeepFinder software, using a clustering radius of 10 voxels, to extract both the coordinates and size of these segments (count of the number of voxels of each segment), with and without the application of a mask (specifically, a cytosol mask sourced from the dataset, utilized to eliminate false positives outside the mask). We compare the extracted coordinates to the expert-validated annotations provided by the dataset at different levels of segment sizes used as a threshold (i.e., segments smaller than the threshold were removed). For this comparison, we maintained the criterion that two annotations—one from the output of the deep learning model and the other from the expert annotations—were considered to target the same particle if their spatial distance was within 10 voxels (a value consistent with the methodology of a previous study analyzed the same dataset ^20^). An example of a tomogram, its corresponding segmentation map, and the coordinates set at a threshold that removed smaller objects compared to expert annotations are presented in Fig. 3. To evaluate the performance of the method, we computed the overall Recall and Precision with and without masking (refer to the Methods section for more details on the assessment metrics). The point where Precision and Recall curves intersect was utilized to determine the F_1_ score per tomogram for the dataset (shown in Fig. 4A). The results, further illustrated in Fig. 4I-J, reveal that Template Learning used to train DeepFinder exclusively on simulations, achieved state-of-the-art performance, outperforming template matching and previous deep learning methods trained solely on annotations derived from the same experimental dataset based on the median F_1_ score (compared to values reported in ^20^).

**Fig. 3:**
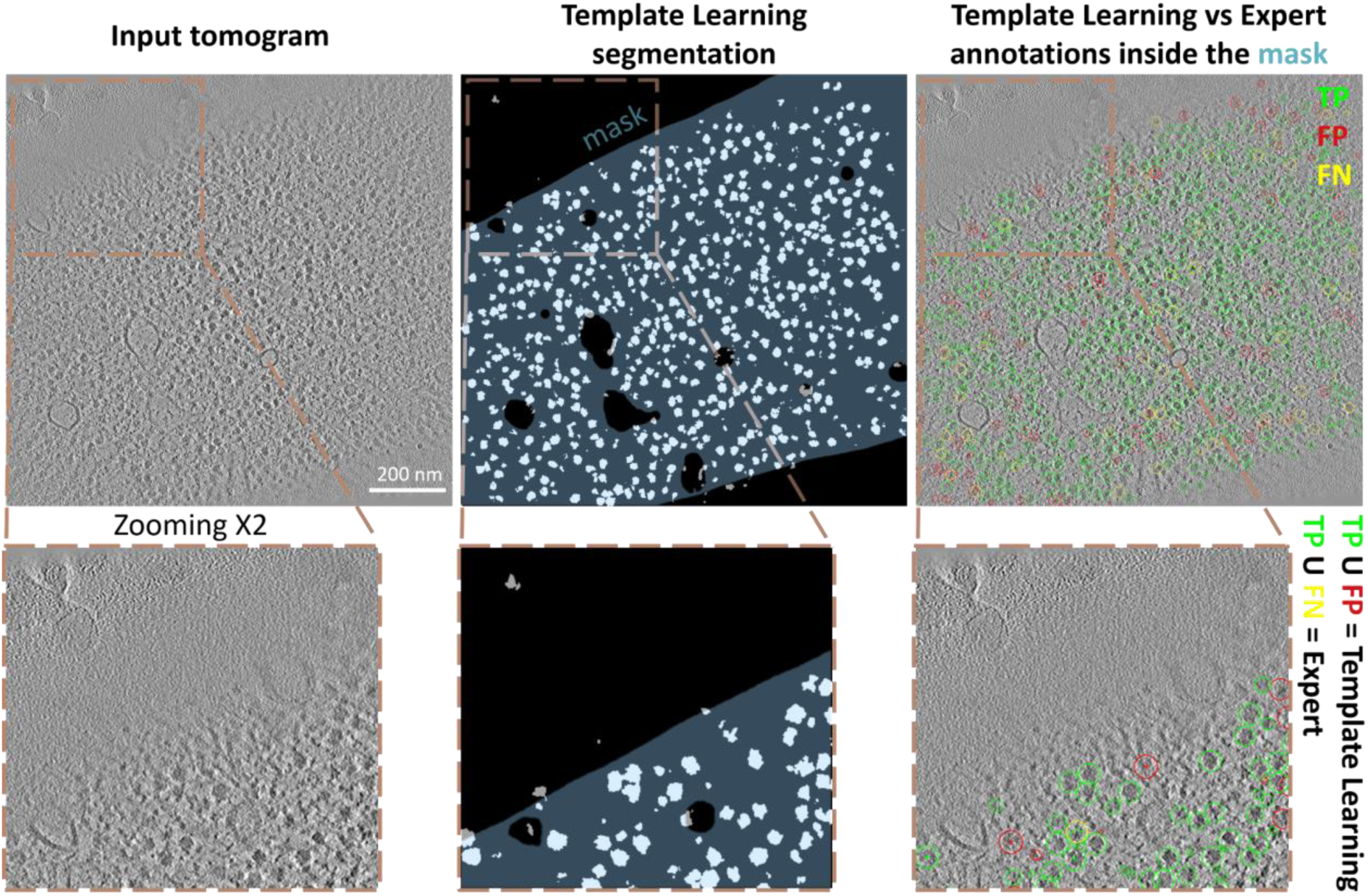
Template Learning trains DeepFinder to single molecule precision on segmenting and annotating ribosomes *in situ*. Left: Display of a 2D central slice of a VPP tomogram. Middle: Segmentation of ribosome from the Template Learning workflow shown in black and white, with a 50% transparent overlay of the mask. Right: resulting annotations within the mask, obtained by converting segmentations to annotations using the MeanShift algorithm (provided by the DeepFinder ^19^ software) using a clustering radius of 10 voxels, compared to expert annotations. Comparison includes True Positives (TP), False Positives (FP), and False Negatives (FN) relative to expert annotations. The tomogram, mask and expert annotations are sourced from the EMPIAR-10988 dataset.

**Fig. 4:**
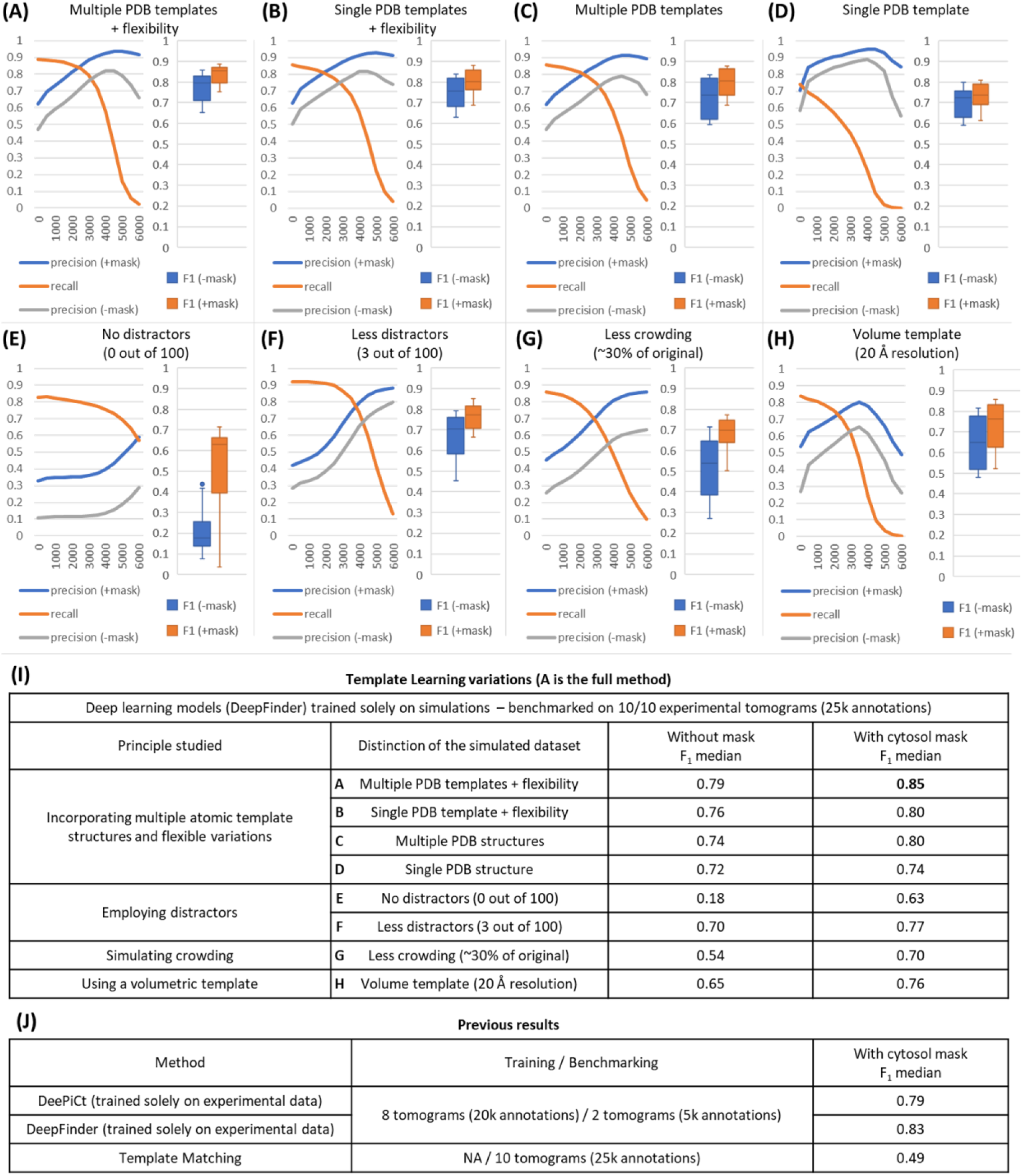
DeepFinder trained on Template Learning simulations only outperforms previous techniques for ribosome annotations in cryo-electron tomograms. **A-H** Performance measures for 8 variations of Template Learning settings. The curves depict the overall Precision and Recall against the volume of the segmented region (horizontal axis). Precision is evaluated with and without masking (±mask). Boxplots show the F_1_ score per tomogram at the threshold corresponding to the highest overall F_1_ score (often coinciding with the intersection of the Precision and Recall curves). **I** Table listing the differences between the experiments in **A-H** comparing their results based on their median F_1_ scores (bold value is the highest among proposed and previous techniques). **J** Table of previous results with their reported performance based on the findings in _20_. Comparison of segmentations resulting from the different experiments on an example tomogram are shown in Extended Data Fig 3.

To reflect on these results, it is essential to highlight that the previously reported findings for DeePiCt and DeepFinder were based on models trained with 8 out of 10 fully annotated tomograms, containing approximately 20,000 ribosome annotations ^20^. Hence, previous evaluations were conducted on 2 out of 10 tomograms, for annotating the remaining 5,000 ribosomes, utilizing three cross-validation splits (in each split, a random set of 8 tomograms was used for training and the remaining 2 tomograms for benchmarking). Additionally, each of the two models underwent a different training strategy to achieve optimal performance. In particular, in the case of DeepFinder, the training encompassed two classes—ribosomes and Fatty Acid Synthase (FAS)—as training solely on ribosomes led to suboptimal performance. On the other hand, DeePiCt was exclusively trained on ribosomes, as unlike DeepFinder, combining ribosomes and FAS during training led to suboptimal performance.

In contrast, the DeepFinder model trained solely on the simulations generated from the Template Learning workflow, not fine-tuned on any annotated experimental data, was benchmarked on the complete dataset (i.e., all 10 tomograms). Hence, despite the relatively modest increase in the F_1_ score for the Template Learning method (0.85, compared to the previous best of 0.83), it stands as the first demonstration that deep learning can be trained effectively for picking a target structure, starting from only prior templates, and domain-randomized simulations. Also, a model of the same network architecture (DeepFinder) trained only on simulations outperforms its counterpart trained only on annotated experimental datasets for cryo-ET particle picking.

### Template Learning variations

We introduced several novel concepts in the Template Learning data simulation workflow, demonstrating that, when combined, they achieve state-of-the-art performance for supervised deep learning in annotating ribosomes within *in situ* cryo-ET tomograms. To study the significance of each proposed concept and explore the potential for integrating various modifications, we conducted additional experiments, which we have organized into the following four categories:

### Incorporating multiple atomic template structures and flexible variations

In three experiments, we progressively reduced the structural variability of input templates while keeping the remaining parameters of the method unchanged. Our motivation behind this series of experiments is to guide potential users regarding the outcomes of employing multiple template structures and integrating flexible variations. These experiments can be particularly useful to future studies, recognizing that, for certain biomolecules, the presence of multiple structures and the feasibility of simulating molecular mechanics may vary.

In one experiment, we retained from the templates a single structure and its flexible variations. In a second experiment, we retained the six structures while excluding the flexible variations. Finally, in a third experiment, we retained only a single structure, eliminating other structures and all flexible variations. The trained deep learning models resulting from the Template Learning workflows of these three experiments were used to analyze the dataset as presented previously.

Fig. 4(B-D, I) depicts the Recall-Precision curves and F_1_ scores for the three experiments. Notably, employing a single structure with flexible variations yielded comparable results to employing multiple structures without flexible variations. We can infer from these findings that introducing artificial flexible variations can serve as an effective compensatory mechanism for the absence of multiple PDBs. This becomes particularly relevant in studies where only a limited number of atomic structures, or even just a single structure, is publicly accessible. Both approaches (i.e., a single PDB with flexibility and multiple PDBs without flexibility) showed a slight adverse impact on the median F_1_ score of the aforementioned typical template learning workflow, of approximately 5%. However, they still demonstrated comparable performance to previously reported results from supervised deep learning on annotated experimental data (compare Fig. 4I/B-C and Fig. 4J).

In contrast, employing a single template without any flexible variations yielded a stricter model characterized by higher Precision and lower Recall—translating to fewer false positives at the cost of more false negatives (see Extended Data Fig.3 for the output segmentation maps). Overall, the use of a single PDB template led to a relative reduction in the median F_1_ score by approximately 10%. Nevertheless, it maintains a significant advantage over traditional template matching (compare Fig. 4I/D and Fig. 4J/Template matching).

### Incorporating distractors

In contrast to our proposed Template Learning method, previous studies did not emphasize the incorporation of a background combining a comprehensive array of dissimilar distractors. In ^48^, no distractors were employed in the data simulation to train deep learning models on subtomogram segmentation. Two recent studies ^27,40^ utilized a small set of objects, serving as analogs to distractors in the simulation of data for training deep neural networks in regression tasks, including signal restoration and segmentation. These objects included gold fiducials, actin bundles, vesicles, and randomly placed small spheres that were relatively sparsely distributed in the volume. Our strategy is principally different; it utilizes other real biomolecules as distractors for a simple approximation of the realistic crowding of biomolecular environments.

To investigate the contribution of employing as many as 100 dissimilar distractors to annotate only a single target structure on the state-of-the-art performance of Template Learning, we conducted two additional experiments on distractors while keeping other method parameters unchanged. In one experiment, we removed all distractors from the simulated tomograms, following a methodology similar to that outlined in ^48^. Fig. 4(E, I) depicts the Recall-Precision curves and F_1_ scores for this experiment, revealing that removing distractors from the simulated data results in hallucinations in the output segmentations leading to a high rate of false positives (refer to Extended Data Fig.3).

In a subsequent experiment, we explored if there are benefits of using many distractors compared to a limited number, similar to previous approaches ^27,40^. Subsequently, we reintroduced three distractors—small (PDB 1S3X), medium (PDB 5A20), and large (PDB 6UP6) proteins—while maintaining the previously established volumetric density ratio balance between distractors and templates in the simulated tomograms. Fig. 4(F, I) depicts the Recall-Precision curves and F_1_ scores for this experiment. These results show that limiting the variability of distractors had a serious adverse impact on the picking Precision.

These experiments suggest that exposing the deep learning model to a diverse range of unwanted structures (i.e., distractors) alongside the target structures (i.e., templates) is essential for the best performance.

### Simulating crowding

To investigate the contribution of simulating high crowding, (i.e., using the Tetris algorithm introduced in this work) to the state-of-the-art performance of Template Learning, we aimed to design an experiment involving crowding reduction. Consequently, we continued our investigation of the crowding reduction impact on Template Learning by adapting the Tetris algorithm to systematically generate crowding levels aiming for 25-35% crowding relative to the original settings while keeping all other parameters unchanged (to see how crowding can be controlled in the Tetris algorithm, see the corresponding Methods section).

Fig. 4(G, I) depicts the Recall-Precision curves and F_1_ scores for this experiment, revealing that the deep learning model trained on a simulated dataset with less crowding resulted in a 20-25% decrease in the median F_1_ score.

### Using a volumetric template

To the best of our knowledge, this work represents the first effort to train deep learning models to utilize fully atomic structures and physics-based simulations as alternatives to volume templates in traditional cryo-ET template matching. An advantage of employing fully atomic structures is the utilization of rapid molecular mechanics simulations using NMA. Particularly, our strategy involves coarse-graining the structures before NMA, and interpolating the resulting motions to the original fully-atomic model (refer to the NMA section in Methods for further details).

However, using volumetric templates, in the context of particle picking, can be advantageous in specific scenarios, particularly when an initial structure can be derived from the dataset. For instance, a subtomogram average at low resolution derived from a partially annotated dataset can serve as an initial template to annotate more particles in the dataset, enabling the generation of a higher-resolution average.

Consequently, we explored a method to incorporate a volume as a template in the physics-based simulation, rather than relying solely on atomic structures. Existing literature demonstrates the feasibility of generating pseudoatomic models from volumes, primarily by estimating the volume using a set of 3D Gaussian functions ^49,50^. The mean and standard deviations of these 3D Gaussians are calculated in a way that the pseudoatomic model, when converted back to a volume (i.e., via summation of the contribution of each 3D Gaussian to each voxel), estimates the volume with low error (e.g., 5%). However, advanced cryo-ET physics-based simulators (e.g., Parakeet, used in this work), do make use of the characteristics of these 3D Gaussians in the simulations, instead, they use electron-atom interactions to model the image formation in the TEM. Meaning, pseudoatomic structures based on previous works do not generate the expected contrast when used in such physics-based simulators.

Hence, we devised a two-stage algorithm for real-time conversion of volumes into pseudoatomic models that allows tuning the contrast by substituting some pseudoatoms with actual phosphorus atoms (see Extended Data Fig.4 for illustration and Methods for details).

To proceed with investigating the effect of using a volume as a template for ribosome annotations, we employed the atomic structure of a ribosome (PDB 4UG0) to generate a volume at 2 nm resolution using the Eman2 ^46^ pdb2mrc software. It is noteworthy that in the study ^20^ where the dataset (EMPIAR-10988) was published, a sub-nanometer subtomogram average was obtained. Here, we explored a hypothetical scenario where a 2 nm resolution initial average is obtained by averaging a partially annotated subset of the original dataset and investigated if the Template Learning workflow can employ this structure for annotating more ribosomes. Subsequently, we transformed this volume into a pseudoatomic structure, following the steps outlined in the corresponding Methods section. As part of this conversion process, we introduced a substitution, replacing one-third of the pseudoatoms with phosphorus atoms to approximate the contrast of ribosomes observed in experimental data (Extended Data Fig.4).

We employed this pseudoatomic model to generate simulations using the Template Learning workflow, by replacing the atomic structures that were previously used as input, keeping the remaining parameters unchanged.

Fig. 4(H, I) depicts the Recall-Precision curves and F_1_ scores for this experiment, indicating that the deep learning model trained on a simulated dataset with the mentioned pseudoatomic structure exhibits a lower Precision and a higher Recall compared to training with an actual atomic structure. We interpret these findings suggesting that lower resolution template structure (2 nm instead of atomic resolution) removed certain details from the simulation, resulting in a deep learning model that is more permissive but less precise in its predictions. Nonetheless, the F_1_ score of this experiment shows a 27% increase relative to traditional template matching, despite that both methods used a similar prior (i.e., a single volume ribosome template).

#### Adapting Template Learning to different domains

In the preceding section, we benchmarked Template Learning for annotating ribosomes in a close-to-focus VPP cryo-ET dataset *in situ*. Hence, the previous Template Learning workflow for ribosome annotation involved simulating data using VPP close-to-focus, consistent with the data analyzed.

In addition to VPP data, the dataset (EMPIAR-10988) contains an additional set of 10 tomograms acquired with defocus (DEF tomograms) and without VPP. In the original study ^20^, the DeePiCt deep learning model was trained on the experimental VPP tomograms and subsequently evaluated on DEF tomograms pre-processed with spectrum matching (SM). SM enhanced the contrast of the DEF dataset to resemble that of the VPP dataset, achieved by extracting a target spectrum from a VPP tomogram and applying it to the DEF tomograms (see Extended Data Fig.5). This approach was referred to as the DeePiCt “cross-domain” experiment.

We conducted two experiments to explore the impact of the data domain on Template Learning. In the first experiment, we set up a Template Learning workflow targeting the DEF domain. We accomplished this by generating a simulated dataset using a procedure identical to the one outlined in the preceding section except for excluding VPP simulations and employing a different range of defocus values (refer to Extended Data Tab.1 for details). This simulated dataset was used to train and benchmark DeepFinder on annotating ribosomes within the DEF tomograms, without undergoing SM preprocessing. In the second experiment, we utilized the DeepFinder model trained on Template Learning VPP simulations to annotate ribosomes within the DEF dataset that had undergone SM preprocessing.

Fig. 5 depicts the Recall-Precision curves and F_1_ scores for the two aforementioned experiments, compared to the previously reported result of the DeePiCt cross-domain experiment. Both of our experiments yield comparable outcomes, outperforming the DeePiCt cross-domain experiment by more than 10% on the median F_1_ score. Notably, the Template Learning-trained model for DEF data exhibited a higher Precision curve and ultimately achieved a higher F_1_ score, particularly when no mask (cytosol mask) was applied.

**Fig. 5:**
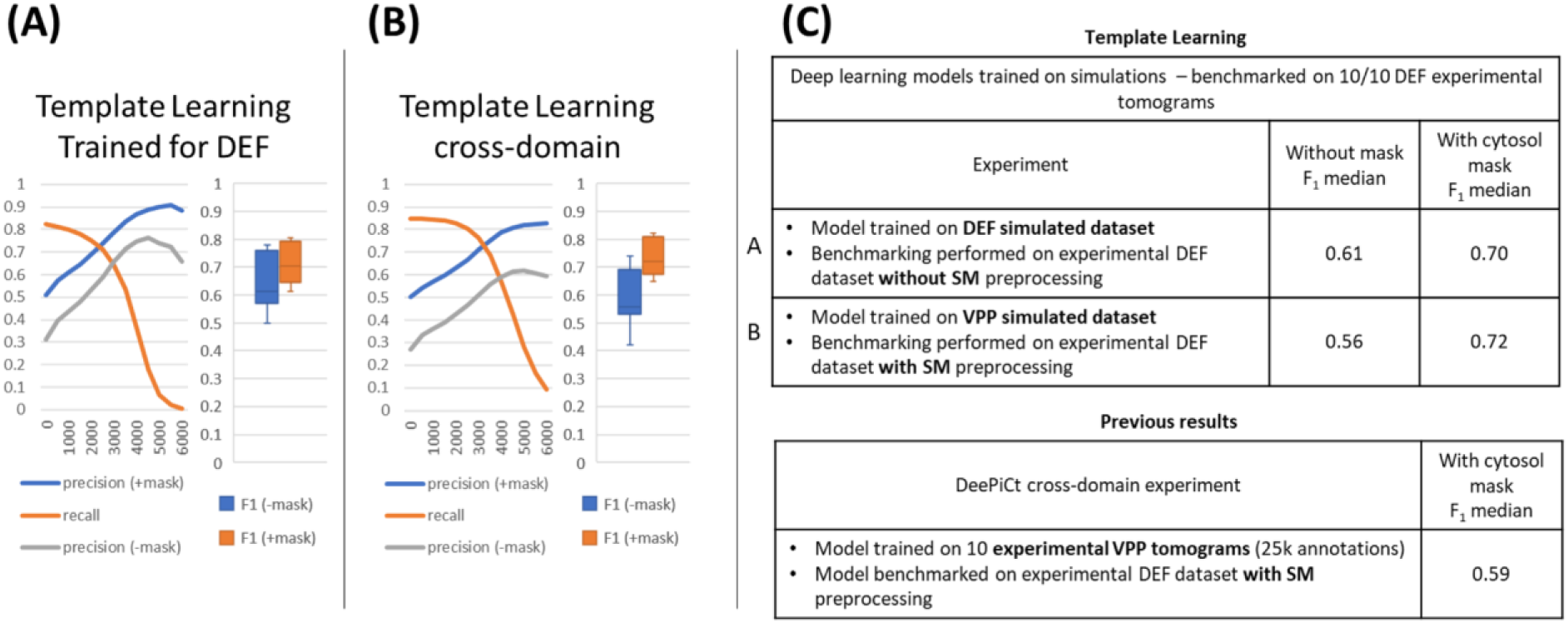
Template Learning does not necessitate data preprocessing for DEF tomograms. Performance benchmarking and comparative analysis of Template Learning workflow applied to ribosome annotations within the S. pombe dataset (EMPIAR-10988, 10 DEF tomograms). **A-B** The curves depict the overall Precision and Recall against the volume of the segmented region (horizontal axis). Precision is evaluated with and without cytosol masking (±mask). Boxplots show the F_1_ score per tomogram at the threshold corresponding to the highest overall F_1_ score (coinciding with the intersection of the Precision and Recall curves). **C** On the top, a table listing the differences between the experiments in **A-B**, comparing their results based on their median F_1_ scores. On the bottom, previously reported performance based on the findings in ^20^.

The first experiment shows that the Template Learning workflow can be directly adapted to the DEF domain avoiding the need for data preprocessing such as SM. The second experiment shows that if a model trained on the Template Learning simulations of the VPP domain exists, it can be reused in the cross-domain (i.e., the DEF domain) after SM, and achieves a similar performance without repeating the entire Template Learning workflow.

### Fine-tuning a deep learning model trained on Template Learning simulations using experimental data annotations

The literature on domain randomization indicates that models pre-trained on simulations and subsequently fine-tuned on real-world data outperform models trained exclusively on real-world data ^35^. This approach can be particularly valuable to cryo-ET particle picking studies that involve challenging-to-annotate molecules, or when training on domain randomization simulations alone does not yield satisfactory results.

In the same dataset used for benchmarking Template Learning on ribosomes *in situ* (EMPIAR-10988), another molecule, FAS, is annotated. Previous attempts based on template matching and supervised deep learning on experimental datasets have reported challenges in annotating FAS, attributed to its distinctive shell-like structure, where particles have lower SNR compared to ribosomes ^20^.

In the following, we present the benchmarking of Template Learning combined with DeepFinder, initially trained exclusively on a simulated dataset, on FAS annotation in EMPIAR-10988 tomograms. Subsequently, we gradually introduce fine-tuning based on the experimental data annotations to assess whether performance can be improved.

To establish a Template Learning workflow for FAS, we utilized two PDB structures (PDB IDs: 4V59 and 6QL5). By using NMA, we generated 80 flexible variations for each structure (refer to the NMA section in the Methods for details). We kept the remaining parameters for simulated data generation consistent with those aforementioned for the ribosome study, except for training the models on sphere segmentations due to the shell-like structure of FAS.

The F_1_ score results of the simulation-trained DeepFinder model for FAS annotations are presented in Fig. 7. The results show that on the VPP dataset, the DeepFinder model trained on the Template Learning workflow outperformed that trained exclusively on experimental data. The results on the DEF dataset gave an F_1_ score ranging up to 22%, outperforming previous attempts that failed to pick any particle, based on the results reported in ^20^. However, these results can still benefit from improvements compared to previously obtained results with DeePiCt trained on annotated experimental data.

We proceeded to fine-tune the simulations-trained DeepFinder model progressively using annotations from experimental data. In the following experiment, we fine-tuned it using annotations from 2 VPP tomograms containing approximately 150 FAS annotations. We kept all the DeepFinder training parameters to default (listed in Extended Data Tab.2), except for reducing the number of steps and epochs to 10 steps for 10 epochs (in place of 100 steps for 100 epochs) to avoid overfitting (since the number of training data examples is low). We performed three cross-validation experiments (i.e., in each experiment, the data is randomly split into 2 tomograms for training and 8 for benchmarking). The results presented in Fig. 6 show a significant increase in the median F_1_ score compared to training on simulated data only.

**Fig. 6:**
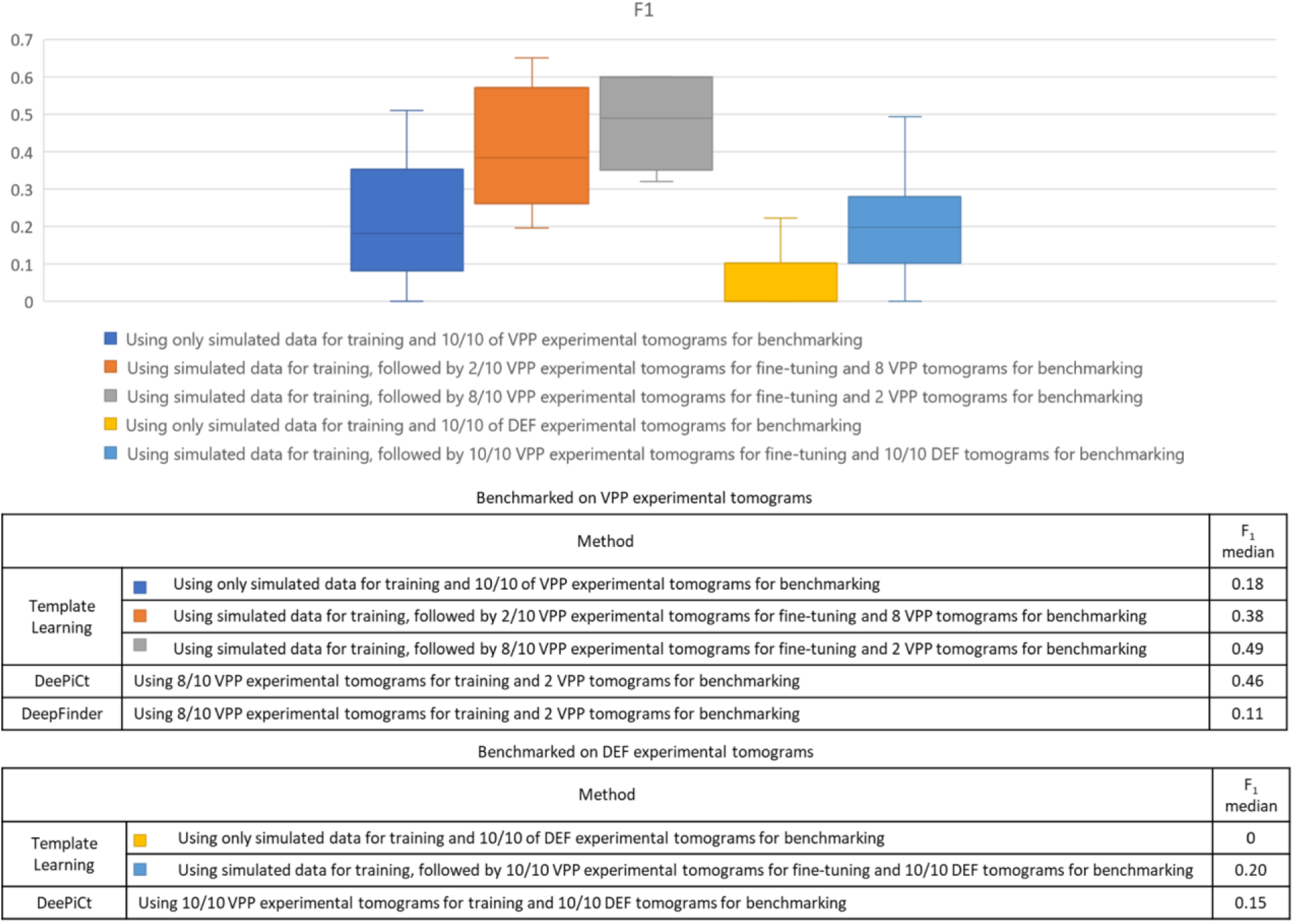
DeepFinder can be pre-trained on Template Learning simulations and fine-tuned on experimental data annotations. Performance benchmarking and comparative analysis of Template Learning applied to FAS annotation within the S. pombe dataset (EMPIAR-10988, 10 DEF tomograms). The results show the performance of the Template Learning workflow used for training DeepFinder on simulations, followed by different levels of fine-tuning on experimental data, compared to results based on the findings in ^20^.

In a subsequent experiment, we fine-tuned the simulation-trained DeepFinder model on 8 VPP tomograms containing approximately 600 FAS annotations and inferred the model on the remaining 2 tomograms, using 3 cross-validation splits. Again, we kept all the DeepFinder training parameters to default, except for the number of steps and epochs, keeping 10 steps for 30 epochs (again, to prevent overfitting). The results presented in Fig. 6 show a significant increase in the median F_1_ score, outperforming previous supervised deep learning methods trained solely on experimental data.

Finally, we performed a cross-domain experiment, where we fine-tuned the simulation-trained DeepFinder model on the VPP dataset, and applied this model to annotate FAS in the DEF dataset after SM preprocessing. Consistently with previous findings, the results presented in Fig. 6 show that training on simulations and fine-tuning on experimental data outperforms training only on experimental data.

### Template Learning offers higher Precision and better orientational isotropy than template matching on nucleosome picking

In cryo-ET data processing, localizing a target biomolecule in a new dataset is a common objective. The application of supervised deep learning to this task requires an initial training dataset of particles of interest extracted directly from the new data. Such a dataset is usually in the order of thousands of particles in different orientations, where more particles are needed for better performance. Manually annotating these particles is time-consuming, making template matching followed by manual elimination of obvious false positives and further curation through subtomogram averaging and classification the most common approach ^19,20^. The principle of Template learning offers a time and computing-efficient alternative to this complex procedure by annotating new data directly in a single step using a model trained on synthetic data.

In this section, we explore the efficiency of Template Learning for nucleosome annotation within a new cryo-ET dataset of partially decondensed mitotic chromosomes *in vitro* (refer to the Method section for sample preparation details) and compare it with template matching-based annotation routine.

A tomogram central slice of our data is shown in Fig. 8A. Despite this dataset being isolated chromosomes *in vitro*, it contains an abundance of structures other than nucleosomes that can be false positives which are: DNA linkers, gold nanoparticles, percoll, and the non-histone components of chromatin.

### Template Learning for picking nucleosomes

To establish a Template Learning workflow, we utilized six nucleosome template structures with PDB IDs: 2PYO, 7KBE, 7PEX, 7PEY, 7XZY, and 7Y00. By applying NMA, we generated 100 flexible variations for each structure (more details can be found in the NMA section of Methods). The remaining parameters of the Template Learning workflow were configured similarly to those used for the workflow established for ribosome and FAS annotation, with a few notable exceptions explained below.

Firstly, recognizing the smaller size of nucleosomes compared to ribosomes, we adjusted the frequency of appearance of nucleosome templates to one nucleosome for every two distractors (in contrast to one ribosome for every five distractors). This adjustment aimed to maintain a balanced volumetric density ratio between distractors and templates in the simulated tomograms.

Secondly, recognizing that nucleosomes have fewer atoms than ribosomes, we observed increased speed in the execution of the physics-based simulator (i.e., Parakeet). This allowed for the generation of larger tomograms for training, all within the same runtime for the ribosome study (see Software requirements section for details). In particular, the tomogram size generated in the Tetris algorithm here was set to 256x256x64 (for a pixel size of 16 Å), in contrast to 192x192x64 used for ribosome simulations.

Lastly, we adjusted the pixel size of the simulated data to 8 Å through binning, closely approximating the pixel size of our experimental tomogram, which had been previously binned four times before annotation (the unbinned pixel size was 2.075 Å).

Consequently, we trained DeepFinder on the resulting Template Learning simulations and applied it to segment our data. The corresponding segmentation map (score map), is shown in Fig. 7B. Upon visual inspection, it is evident that that model has high Precision in localizing nucleosomes. Subsequently, we employed the MeanShift algorithm from the DeepFinder software to extract the corresponding annotations for the nucleosomes, utilizing a clustering radius of 6 pixels, approximately equivalent to the nucleosome’s radius at this pixel size. We excluded particles close to the edges of the tomogram, the air-water interface, and the sample-carbon interface.

**Fig. 7:**
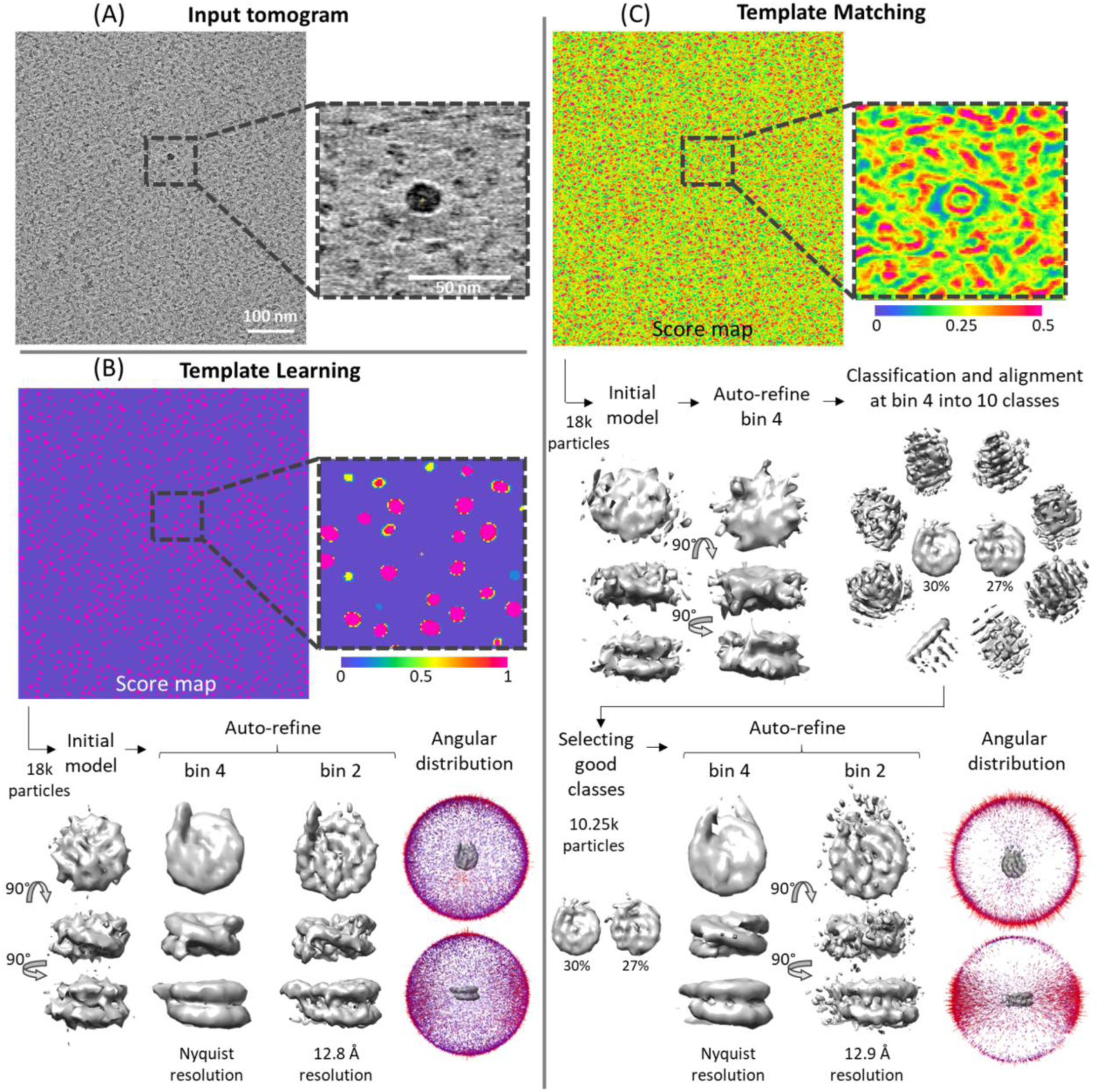
Template Learning outperforms traditional template matching in Precision and orientational isotropy in annotating nucleosomes. **A** Central slice and zoomed-in 10-slice average of a cryo-ET tomogram of partially decondensed mitotic chromosomes *in vitro*. **B-C** The results of applying Template Learning (6 PDB structures as input, other parameters kept default) and traditional template matching (using PyTom with a nucleosome template) for annotating nucleosomes within the tomogram in A, followed by reference-free subtomogram averaging in Relion V4. **B** Top: score map resulting from training DeepFinder on nucleosome annotations using simulated datasets only from the proposed Template Learning workflow. Below: results of subtomogram averaging using 18k annotations extracted using 6 voxel clustering radius of the MeanShift algorithm (provided by the DeepFinder software). The results show initial model generation, followed by two stages of auto-refinement, and the angular distribution of averaged particles. The refinement has converged without particle curation and the angular distribution of averaged particles is isotropic. **C** Top: score map resulting from traditional template matching. Below: results of subtomogram averaging using the top 18k local maxima from the score map. The first step was an initial model generation, followed by a stage of refinement. The first stage of refinement did not improve the resolution of the initial model, therefore, it was followed by simultaneous classification and alignment into 10 classes. The particles corresponding to nucleosome-like classes (57% of the original 18k particles) were used for two stages of refinement. The results show a slightly lower resolution average than the one obtained using Template Learning and show an anisotropic angular distribution.

Following this process, we obtained 18k annotated particles and proceeded with the reference-free subtomogram averaging procedure using Relion V4.0 ^15^ by generating an initial model from the data, followed by two stages of refinement at binning 4 and binning 2, resulting in resolving the nucleosome structure at 12.8 Å (at 0.143 FSC threshold) resolution shown in Fig. 7B.

The angular distribution of the averaged particles seems isotropic, showcasing that the Template Learning method is not biased to certain orientations.

To further verify the Precision of nucleosome identification, we classified the particles into 10 and 50 classes (presented in Extended Data Fig.6). All class average results showed well-recognizable nucleosome shapes with DNA gyres, with the major differences being in the heterogeneity of the DNA entry-exit segments. We also performed a local resolution assessment using ResMap ^51^ (presented in Extended Data Fig.7) and observed that resolution has ranged from 12-20 Å, with the highest resolution in the DNA close to the core histones, and the lowest resolution around the DNA entry-exit segments.

Our averaging and classification results show that our Template Learning workflow generated precise nucleosome annotations of uniform orientations, facilitating high-throughput subtomogram averaging without the need for multiple rounds of classification to eliminate false-positive particles.

### Template matching for picking nucleosomes

We applied template matching starting from a nucleosome template structure generated from the PDB 2PYO using the default procedure in PyTom ^18^. We then applied the template matching procedure to our new data using an angular sampling of 7° increment (45,123 rotations).

The score map is shown in Fig. 7C. The visual inspection of the score map shows some high signal for some nucleosomes also identified by the Template Learning procedure (Fig. 7A), but also obvious false positive signal for other objects (an example of high false signal for the Percoll particles is shown at the center of the zoomed-in image in Fig. 7C).

To have a meaningful and fair comparison with the Template Learning procedure, we extracted from the same tomographic region the 18k particles showing the highest cross-correlation scores. Unlike the results of Template Learning, a brief visual inspection of the template matching peaks showed 50 obvious false positives corresponding to the 10 nm gold and Percoll particles. The false positives were removed, and data were processed Relion V4.0 ^15^. The reference-free initial model was generated, followed by a stage of refinement at binning 4. Unlike Template Learning, the refinement did not result in a significant improvement of the resolution of the initial model (judged by the shape and the FSC), indicating the presence of a significant ratio of non-nucleosome particles among the annotations. In agreement with this assumption, classification, and alignment into 10 classes at the same binning resulted in only 2 nucleosome-like classes that sum to 10.25k picks (57% of the original picks). The particles of these 2 classes were selected for a further refinement process at binning 4 reached Nyquist resolution, and the further refinement at binning 2 led to 12.9 Å resolution which is comparable to that achieved with Template Learning.

Importantly, the angular distribution of nucleosomes annotated by template matching showed a significant imbalance towards side views compared to round top views (Fig. 7C). Our simulations (Extended Data Fig.8) showed that due to the cylindrical shape of nucleosomes and the missing wedge problem of cryo-ET data, the constrained cross-correlation between a template and the particles is a function of the orientation, where side views show higher cross-correlation than top and oblique views. This problem results in extracting only side views of nucleosomes at an adequate Precision.

The orientation bias of template matching hampers its efficiency for particle detection and is prone to resolution loss in subtomogram averaging. In our case, the resolution of the nucleosome was not affected significantly, because its pseudo-symmetric cylindrical shape allowed the complete 360° orientation range of side views to have cross-correlation sufficient for detection (Fig. 7C). However, this constraint may pose a more significant challenge when annotating particles with asymmetric non-spherical shapes. Our results demonstrate that Template Learning is capable of overcoming this limitation.

## Conclusion

In this study, we introduced Template Learning, a novel approach that blends the simplicity of Template Matching, requiring only template structures as input, with the advanced capabilities of Deep Learning, thereby reducing the dependence on extensive, labor-intensive annotated datasets for supervised training. Our method effectively tackles key challenges in cryo-ET particle picking, such as structural variability, cellular crowdedness, and experimental data variance, through a refined implementation of domain randomization to create simulated data for deep learning training. The efficiency of Template Learning arises from its focus on essential aspects, enabling the well-established DeepFinder model, known for its particle annotation capabilities, to train without the need for highly realistic cellular environment simulations. This approach simplifies the process for users, allowing them to generate effective simulations that train DeepFinder in annotating specific biomolecules in cryo-ET data, using only templates in a streamlined yet customizable simulation process. Template learning can utilize any atomic models, including those generated by artificial intelligence-based tools, as templates. We also confirm the feasibility of relevant template generation from low-resolution cryo-EM densities.

In this paper, we employed a comprehensively annotated *in situ* cryo-ET dataset for ribosomes and FAS, covering defocused cryo-ET and VPP, to validate Template Learning’s efficacy and versatility. We demonstrated how to enhance DeepFinder’s supervised training by initially training on simulations, and then fine-tuning on experimental data. Notably, DeepFinder, when trained solely on simulations, surpassed previously established training on experimental annotations for ribosomes. Furthermore, the combined approach of pre-training on simulations and subsequent fine-tuning on experimental data showed improved performance for FAS.

We evaluated the efficacy of our method for the identification of a known target molecule in tomographic data without any prior annotation, focusing on the case of localization of nucleosomes in isolated chromosomes *in vitro*. Template Learning outperformed the conventional template matching and classification routine typically employed for this type of task both in terms of annotation Precision and orientational isotropy. Notably, all annotations generated by Template Learning could be directly used for achieving high-resolution subtomogram averaging without requiring any manual or classification-based curation.

In future work, we envision adapting our framework to locate structures bound to other biomolecules such as membrane proteins. Another challenge will be to reduce computational issues as physics-based simulations are still resource-intensive.

Conclusively, we believe Template Learning’s versatile and straightforward framework marks it as a timely and potent tool for a broad spectrum of cryo-ET studies, ready to make significant impacts in the field.

## Software requirements and availability

Template Learning code is freeware, open source, GPU and CPU optimized, and fully implemented in Python (https://github.com/MohamadHarastani/TemplateLearning). It provides all the steps that are required to reproduce the results presented in this article and can be straightforwardly applied to other studies. It provides wrappers for ProDy, Eman2 and Parakeet that will be installed during the installation. The software was tested on our workstations (Dell Precision 5820, Intel(R) Xeon(R) W-2145 CPU @ 3.70GHz, 96 GB DDR4 RAM, 2 X NVIDIA RTX A6000 or NVIDIA Quadro RTX 8000). The time required to perform the Template Learning workflow ranged from 1 to 2 days (depending on the number of used GPUs). Installation and user guides are available on GitHub.

## Data availability

The tilt series, tomogram, coordinates, and other metadata necessary to reproduce the nucleosome subtomogram averaging are deposited on EMPIAR via the accession code EMPIAR-11969. The subtomogram averages resulting from the Template Learning and template matching workflows are available on EMDB via the accession codes EMD-19823 and EMD-19825 respectively. Other benchmarking datasets employed in this work are publicly available.

## Author contributions

Investigation, conceptualization, and methodology were done by MH and ME with contributions from all authors. Software programming and computational experiments were done by MH. Sample preparation and data acquisition were done by GP under the supervision of ME. Subtomogram averaging was done by MH and GP under the supervision of ME. Validation and results interpretation were done by all authors. Writing the original draft was done by MH with input from GP. Review and editing were done by MH, ME, and CK. Funding acquisition was done by ME and CK. Project administration was done by ME.

## Methods

### Normal Mode Analysis (NMA)

Our study employed NMA on various atomic structures, prioritizing computational efficiency over precise molecular mechanics simulation, as the primary focus was on training the network on variations of the target biomolecule rather than precisely predicting its conformations. This emphasis on expanding the domain of the synthetic data aimed to capture a broad range of variations, not solely constrained to those witnessed in experimental data. To manage computational demands, we adopted a coarse-grained modeling strategy ^53^, focusing on Carbon Alpha and Phosphorus atoms for NMA and subsequently expanding (interpolating) the modes to the complete atomic structure. Typically, a substantial number (three times the number of atoms in the atomic structure) of normal modes are calculated simultaneously, which can be ordered based on their frequency. Low-frequency normal modes are particularly useful as they represent global movements, while high-frequency normal modes depict local movements. Prior studies have consistently highlighted the significance of these low-frequency, high-collective modes in capturing experimentally observed conformational variabilities ^54,55^.

The user of NMA retains control over the selection of modes and the amplitude of deformation to generate flexible variations, ensuring a balance between simulating variability and maintaining structural integrity. As a guideline, we chose the first 20 low-frequency normal modes for generating random flexible variations. Setting the amplitude range to 150 for ribosomes and 100 for nucleosomes resulted in Root Mean Square Deviation (RMSD) values of approximately 1 Å and 2 Å from the initial structures, respectively. Adjusting this parameter incurs minimal computational costs (a few minutes) by generating variations and assessing the RMSD from the initial structure, for instance using ChimeraX ^52^.

The computational time required for the calculation of NMA varies depending on the size of the input atomic structure. Specifically, for an atomic structure of 200 kDa, such as a nucleosome, the calculation typically takes approximately 1 minute. In contrast, for larger structures like a ribosome weighing around 4.5 MDa, the calculation time extends to about 1 hour when processed on a single CPU (benchmarked on Intel(R) Xeon(R) W-2145 CPU @ 3.70GHz).

In this work, all NMA calculations were performed using ProDy ^56^, a widely recognized open-source Python package for molecular mechanics simulations.

### Tetris algorithm for generating highly crowded samples

The Tetris algorithm (flowchart shown in Fig. 2) utilizes input volumes of the biomolecules, essentially volumetric versions of the PDB templates and distractors, to generate a densely packed sample comprising randomly rotated biomolecule copies. It systematically places one randomly rotated molecule at a time in the closest proximity to previously positioned molecules. The algorithm terminates when a new copy cannot be placed in the sample.

In the initial step, the Tetris algorithm places a molecule at the center of a larger volumetric sample, the size of which is set by the user, establishing the current Tetris output. Subsequent iterations involve randomly rotating a new molecule, followed by its conversion to binary form after applying low-pass filtering and thresholding. While the values for low-pass filtering and the volume threshold are empirical and subject to variation, a standard deviation sigma of 2 for a Gaussian low-pass filter, and a volume threshold of 100 have generally proven effective in our trials (note that the volumes are generated from PDBs using Eman2 ^46^ e2pdb2mrc software). This copy (molecule) is then diluted using a ball-like structural element of user-defined radius. Multiplying the output of the binarized volume by a large positive number (integer infinity in programming) and subtracting the result from the dilated volume, results in an “insertion shell”, which serves as a spatial guide for placing the next molecule. Notably, the radius of the structural element provides direct control over the compactness of the sample.

The correlation between this insertion shell and a binary version of the current output (the binary version is smoothed and thresholded the same way as above) generates a correlation map. White voxels in the correlation map represent potential locations for adding the copy, ensuring the desired intermolecular distance. In contrast, black and gray voxels represent locations where the copy would intersect with previous copies or be too distant from other molecules, respectively. If the correlation map is not entirely negative (stopping criteria), the current copy is added at the index of the maximum entry, corresponding to the location closest to the sample within the desired distance.

During Tetris sample generation, the oriented coordinates used for placing molecules in the volumetric sample are preserved. To generate cryo-ET simulations of this sample, the oriented coordinates are passed to the physical-based simulator (Parakeet).

Generating volumetric samples using the Tetris algorithm with molecules at a resolution of 32 Å and a voxel size of 16 Å^3^, as implemented in our current software, achieves a placement rate of approximately 1000 molecules per minute. The creation of a typical Tetris, with dimensions of 3072 x 3072 x 1024 Å^3^, requires approximately 5 to 6 minutes on a single CPU (benchmarked on Intel(R) Xeon(R) W-2145 CPU @ 3.70GHz) and can accommodate around 5000 molecules. Consequently, the generation of 48 Tetris samples necessary for the tests performed in this study takes approximately 4.5 hours on a single workstation.

### Performance assessment metrics of particle picking

In evaluating how well our deep learning models pick particles, we used the commonly employed metrics in particle localization studies, namely: the Precision, Recall, and F_1_ score. Precision measures the percentage of correct picks out of all the particles selected by the model (True Positives / Positives). Recall measures how many of the actual particles were correctly identified by the model (True Positives / Ground Truth). F_1_ score, being the harmonic mean of Precision and Recall, provides a comprehensive measure of overall model performance.

### Converting volumes to pseudoatoms with contrast tuning

To convert a volume to a pseudoatomic structure that allows physics-based simulations, we devised a two-stage algorithm. In the initial stage, we binarize the volume, similar to the process of generating tight 3D masks in cryo-ET and cryo-EM studies. Subsequently, we assign a pseudoatom to the position of every non-zero voxel, resulting in a point cloud as output. We parse this point cloud into a PDB format, representing each point (i.e., pseudoatom) as a density (DENS line in PDB format). Then, the resulting file is ready to be integrated into the physics-based simulations. However, projections of this structure lack the desired contrast compared to those observed in experimental data. This discrepancy arises from the inherent modeling of electron-atom interactions in physics-based simulators. To address this contrast issue, in the second stage, we substitute a fraction of pseudoatoms with actual atoms, such as phosphorus (e.g., replacing one out of three pseudoatoms with phosphorus atoms). This substitution is not aimed at replicating the precise chemical composition of the structure (a direction that may be explored in future research) but rather to empirically enhance contrast. We determine this conversion ratio (i.e., pseudoatoms to phosphorus) by projecting the pseudoatomic model in 2D and comparing it to the experimental data targeted for analysis.

### Cryo-ET of partially decondensed mitotic chromosomes

#### Chromosome Purification

Chicken DT40 cells were maintained at a density of 8-10 x 10^5^ /ml at 39 ℃ and 5% CO_2._ The cells were synchronized to mitosis using 0.5 µg/ml of nocodazole for 13 hr resulting in a mitotic index of 70 - 90%. The mitotic chromosome isolation was performed using the classic polyamine-EDTA buffer-based method optimized for DT40 ^57^. The synchronized cells were harvested by centrifuging the culture at 1600 g for 5 mins at 4°C. The cells were swollen at room temperature for 5 minutes in a low salt buffer in the presence of polyamines and then lysed using a dounce homogenizer. The lysate was then overlaid on a step sucrose gradient (15%, 60%, and 80% [w/v]) centrifugation. The 60-80% interface was recovered and sedimented on a self-forming percoll gradient in the presence of polyamine. The band containing the chromosomes was recovered and washed to remove the excess of percoll. The isolated chromosomes are stored in Tris-HCl buffer (pH 7.5) containing polyamines in 60% glycerol.

#### Sample preparation and data acquisition

The DT40 chromosomes were vitrified in glow-discharged (operating at 30% power with a gas mixture of 80% Argon:20% Oxygen) 200 copper mesh, Quantifoil Multi Holes grids (Quantifoil Micro Tools GmbH, Germany).

The isolated mitotic chromosomes were exposed to low ionic strength buffer by 100-fold dilution in an aqueous solution containing TEEN buffer (1 mM Triethanolamine:HCl pH 8.5, 0.2 mM Na-EDTA pH 9 and 10 mM NaCl) for 1 hour at 4 ℃. This resulted in swelling of the mitotic chromosomes which were then carefully spun down onto the surface of the grid. 10 nm BSA Gold Tracer (EMS, Hatfield, PA, USA) was added to the grid at a 5:1 ratio (chromosome:gold). The grid was blotted from the carbon side using a teflon sheet and the metal side using a blotting paper respectively. The grid was blotted for 25 seconds with a blot force of 10 and flash-frozen into liquid ethane using Vitrobot Mark IV (FEI) at 4 ℃ and 100 % humidity.

The tilt series was recorded on Titan Krios G3 (Thermo Fisher Scientific) equipped with a Quantum energy filter with a slit width 20eV for higher magnifications and Gatan K2 detector using SerialEM Version 4.1.0 beta ^58^. The images were acquired at a pixel size of 2.075 Å/pixel at 2–4.0 µm defocus at a dose rate of 3.2 e^−^/Å^2^ per image fractionated over 10 frames. A dose symmetric tilt scheme ^59^ was used with a 3-degree increment step and the tilt range was set to ±60 degree using Serial EM.

### Data processing

Individual movies are motion-corrected and averaged to form a tilt series using MotionCor2 ^60^. Gctf was used to perform CTF estimation ^61^. The tilt series were aligned and processed with IMOD^47^. The position of the grid in each image is aligned by tracing 10 nm gold particles as fiducials throughout the tilt series. The aligned tilt series were reconstructed into 3D tomograms with weighted back-projection with simultaneous iterative reconstruction technique (SIRT)-like filter after 4-fold binning.

After picking nucleosomes with our Template Learning trained DeepFinder model, we performed a reference-free subtomogram averaging following the procedure suggested in Relion V4.

## Extended Data

**Extended Data Fig. 1:**
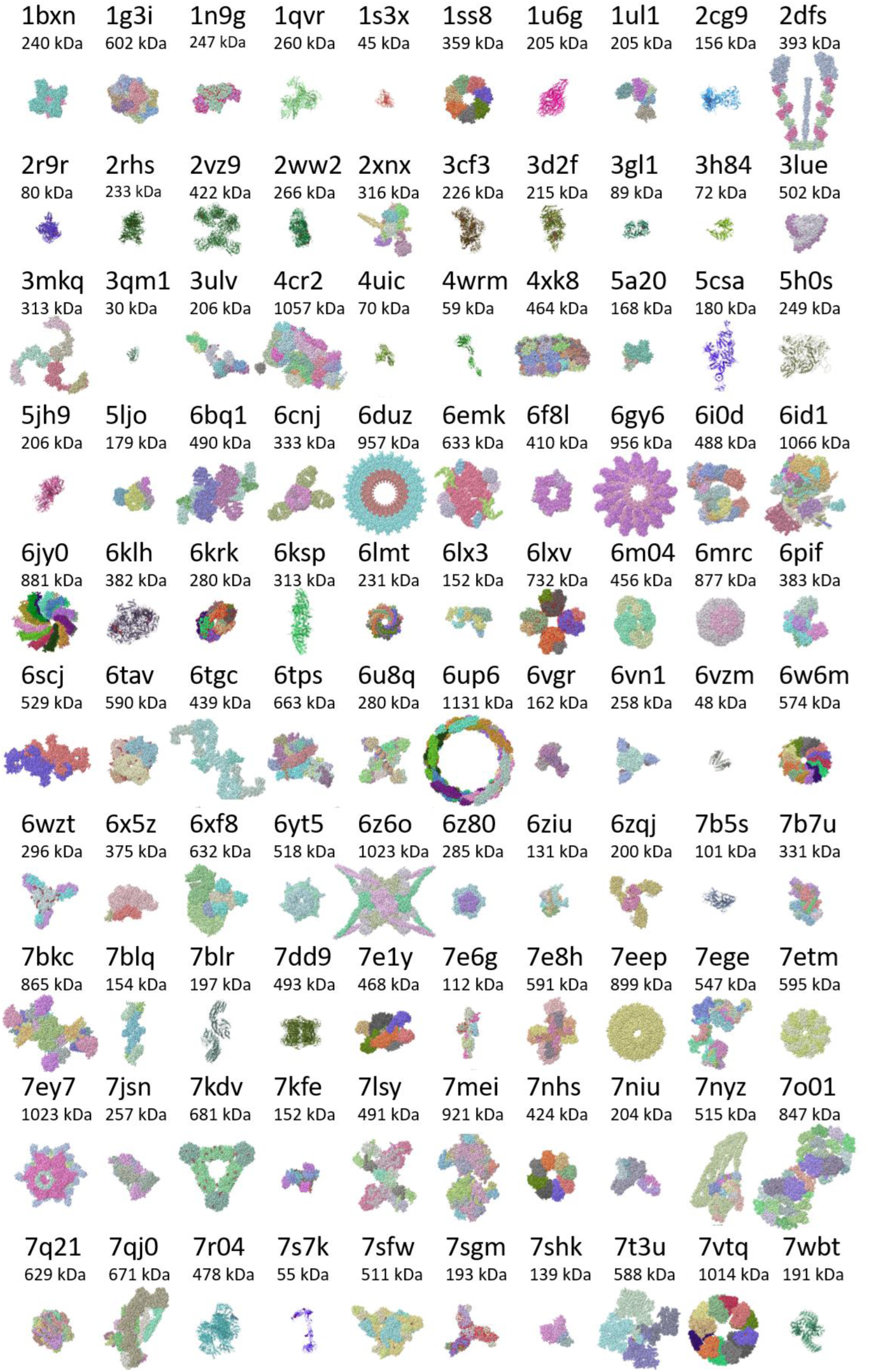
The list of distractors. Template Learning employs a list of 100 dissimilar protein assemblies, termed distractors, used for domain-randomized cryo-ET data simulations. In this figure, the distractors are displayed with corresponding PDB IDs and molecular weights. Display of these structures was done using ChimeraX ^52^.

**Extended Data Fig. 2:**
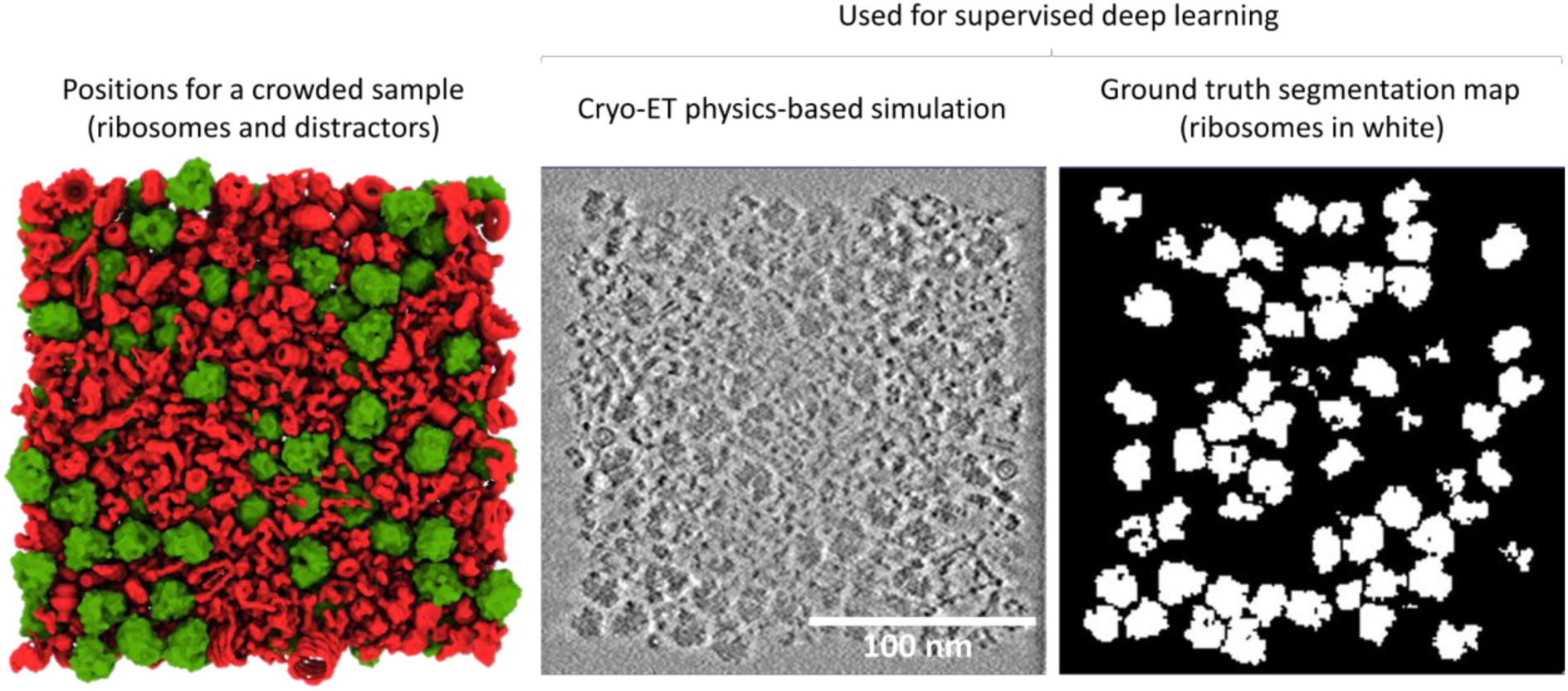
Example of synthetic data based on the Template Learning workflow for supervised deep learning on ribosome annotation. Left: the positions for a crowded sample of size 3072x3072x1024 Å^3^ determined using the Tetris algorithm for ribosomes (in green) and distractors (in red), which are fed to Parakeet (cryo-ET physics-based simulator). Right: a central slice of simulated data and corresponding ribosome segmentations using VPP close-to-focus parameters.

**Extended Data Fig. 3:**
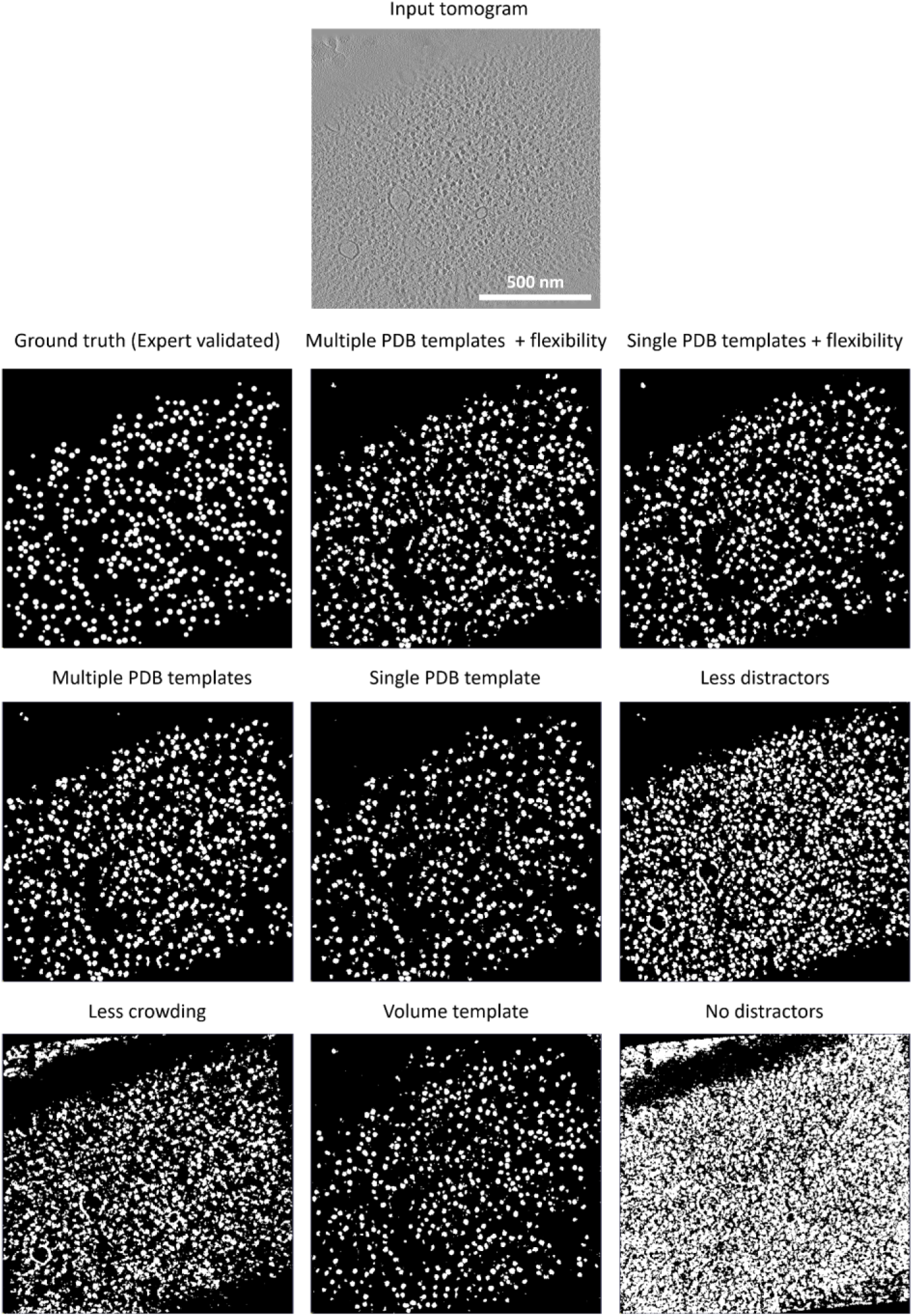
Segmentation maps resulting from different Template Learning variations for ribosome annotations. Central slice of a VPP tomogram from EMPIAR-10988 with its expert-validated (ground truth) ribosome segmentation and the output segmentation maps using the different Template Learning variations presented in **Fig. 4**.

**Extended Data Fig. 4:**
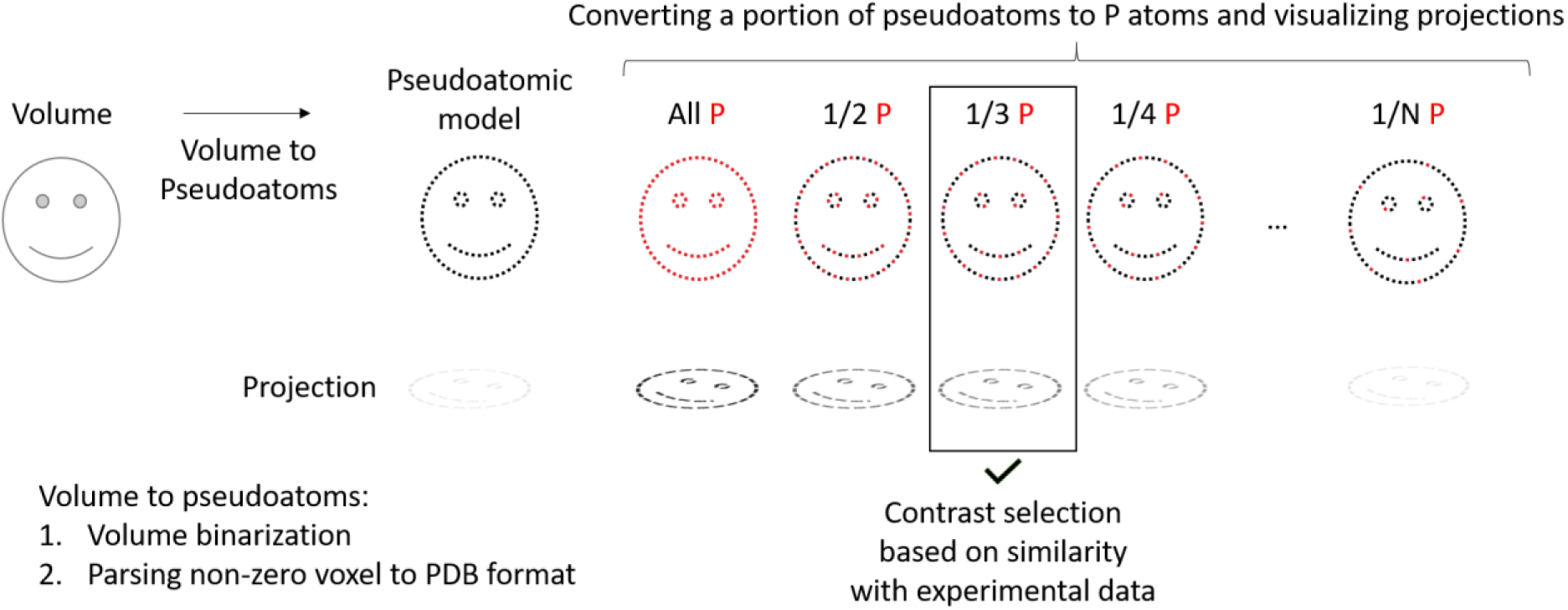
Template Learning can use volumes as templates via volume-to-pseudoatoms conversion. Volume to pseudoatomic model conversion algorithm to produce structures that can be contrast-tuned for usage in physics-based cryo-ET simulators (Parakeet). The method starts by binarizing the volume, a process akin to creating a tight mask through low-pass filtering and thresholding. All non-zero voxels of the binarized volume are parsed into “pseudoatoms”, represented in PDB file format utilizing “DENS” entries as atom types. The generated pseudoatomic structure can be used directly for cryo-ET simulations in Parakeet, but it does not necessarily produce a similar contrast to experimental data as when using fully atomic structures. Therefore, the method allows replacing gradually a portion of the pseudoatoms with phosphorus atoms to emulate more contrasted simulated projections. The optimal proportion is determined by the similarity in contrast, comparing the resulting projections to experimental data. The resultant pseudoatomic model is used for further Template Learning simulations.

**Extended Data Fig. 5:**
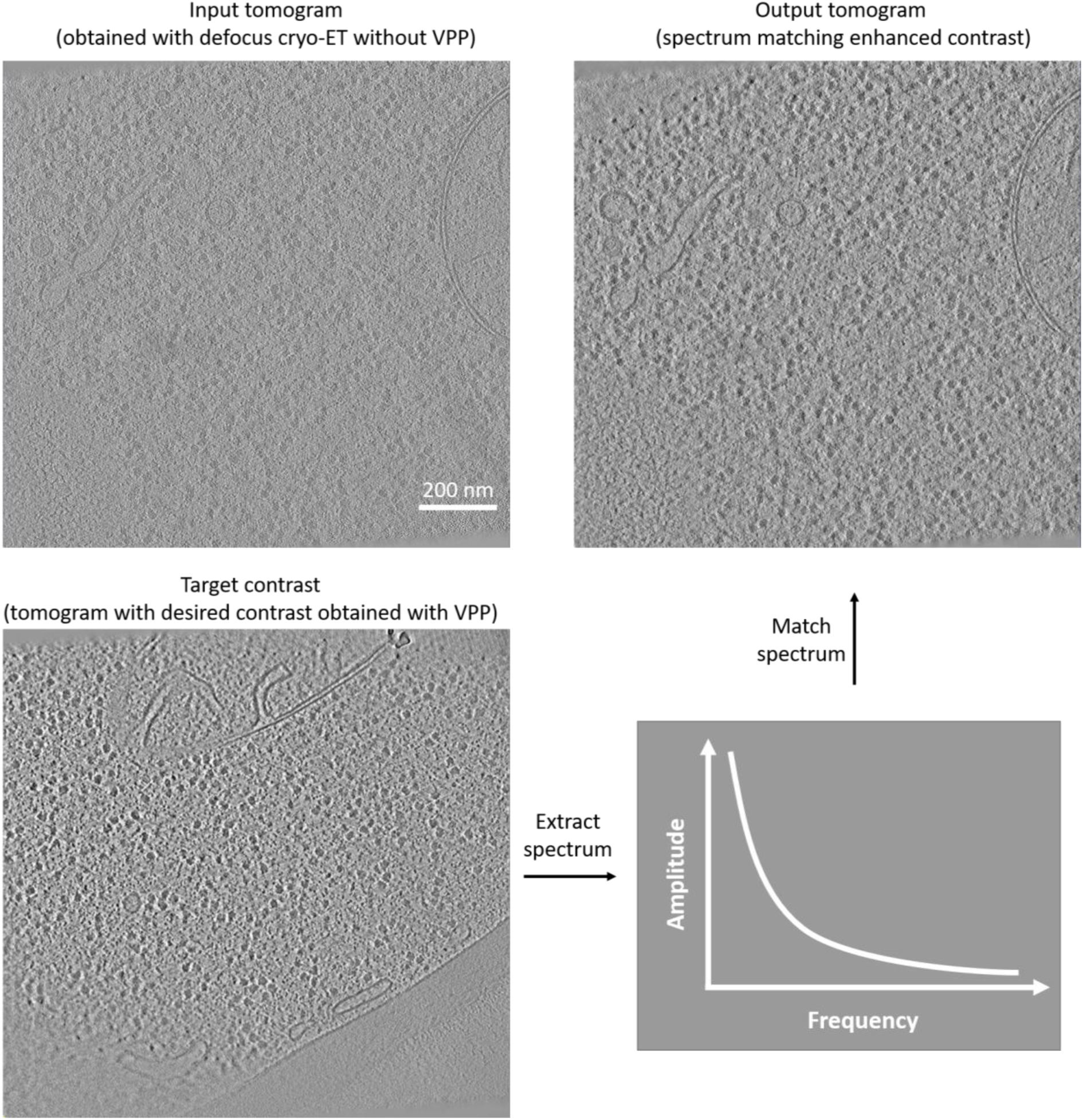
Spectrum matching applied to enhance the contrast of an input tomogram obtained with defocused cryo-ET without VPP. A target spectrum is extracted from a tomogram obtained with VPP, and is applied to enhance the contrast of another tomogram obtained with defocus cryo-ET without VPP, via spectrum matching as proposed in ^20^. Input and target tomograms are sources from EMPIAR-10988

**Extended Data Fig. 6:**
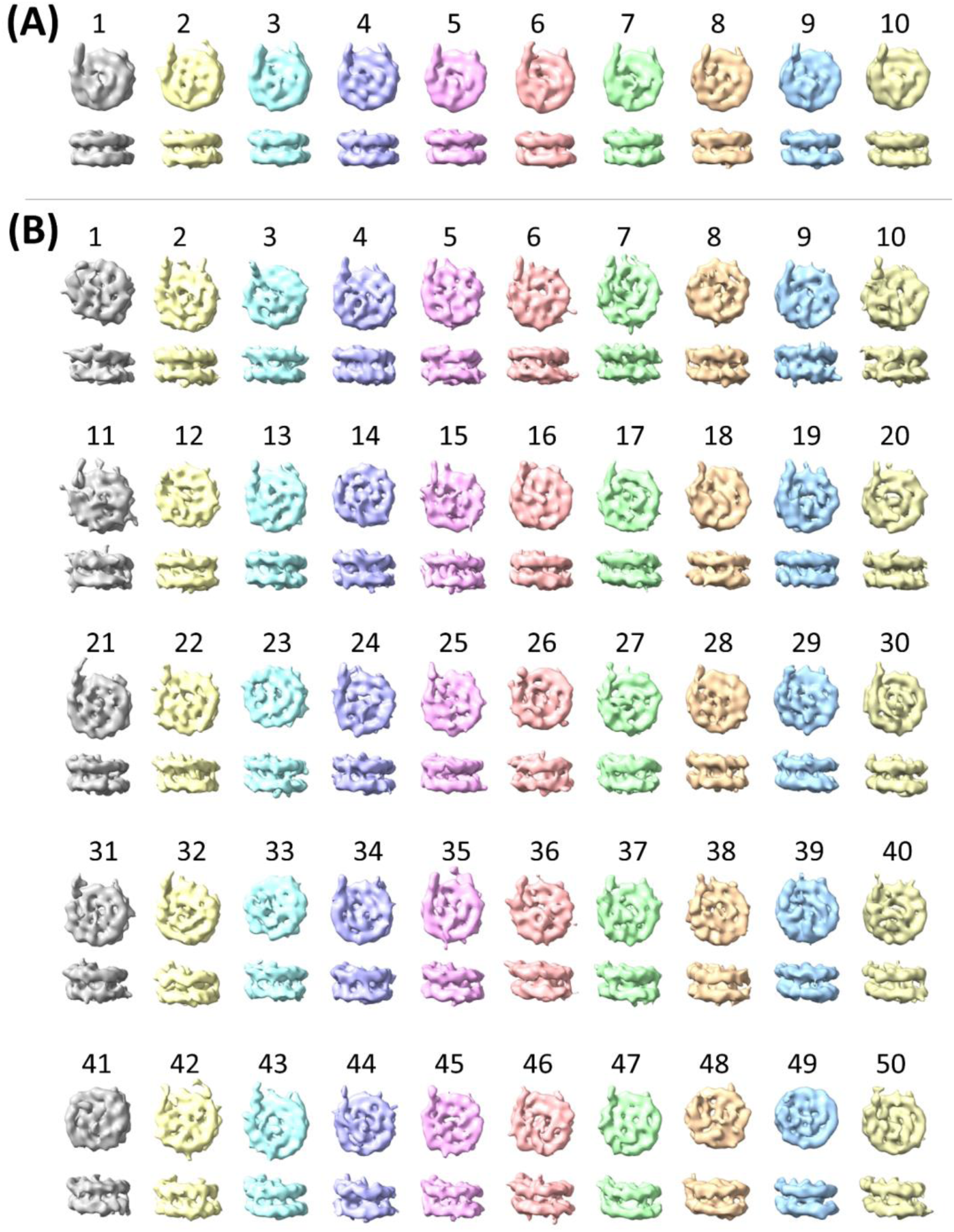
Two experiments of post-alignment classification for nucleosome particles annotated with Template Matching workflow. **A** Classification into 10 classes. **B** Classification into 50 classes. Both classification experiments were done in Relion V4 ^15^.

**Extended Data Fig. 7:**
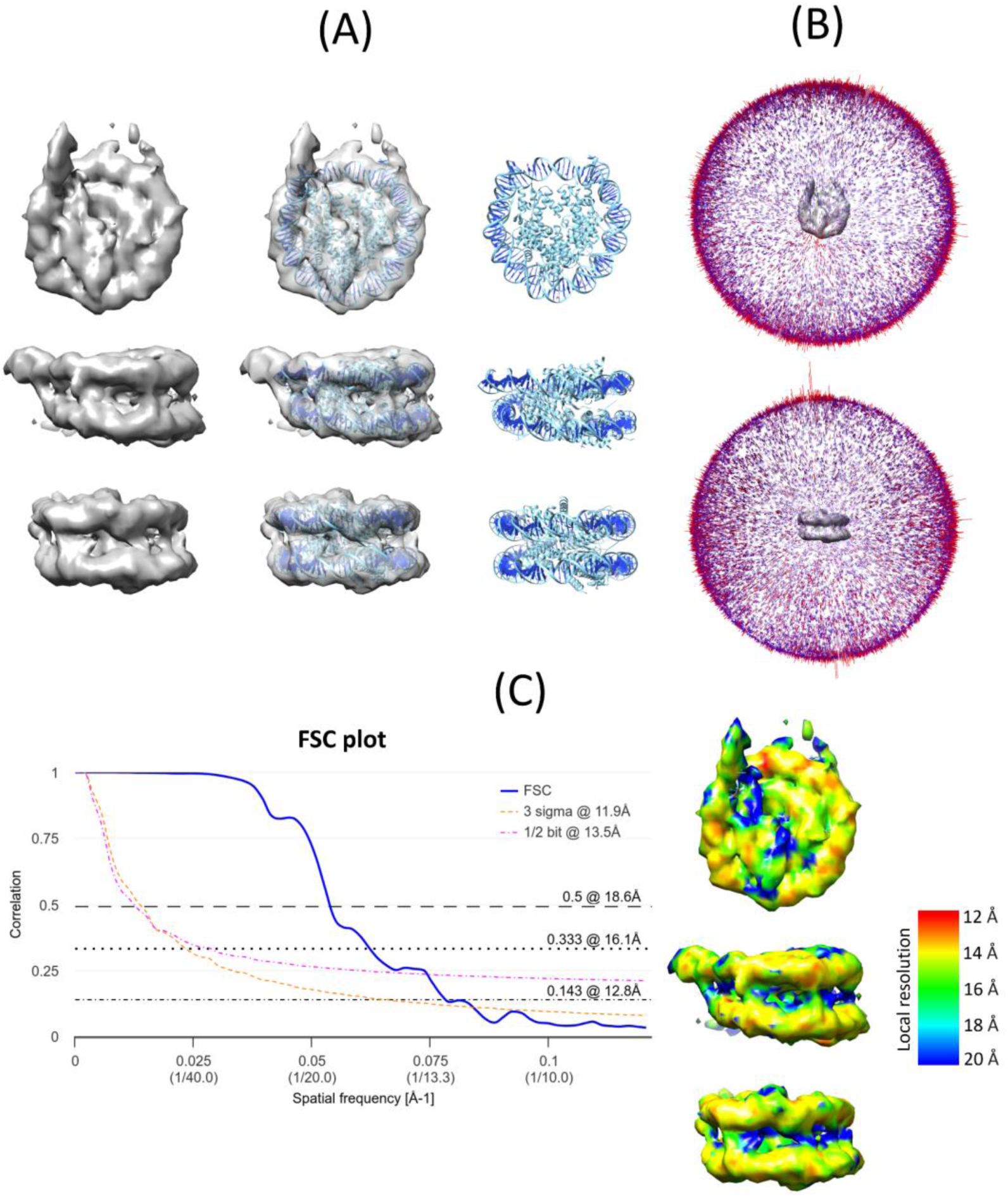
Analysis of the nucleosome subtomogram average obtained via Template Learning annotations. **A** The nucleosome average compared to an available structure (PDB ID: 2CV5). Left: the subtomogram average. Middle: atomic structure docked in the subtomogram average displayed at 50% transparency. Right: atomic structure. **B** Angular distribution of the particles contributing to the global average. **C** Analysis of the resolution of the average. Left: Fourier Shell Correlation curves. Right: local resolution analysis using ResMap ^51^.

**Extended Data Fig. 8:**
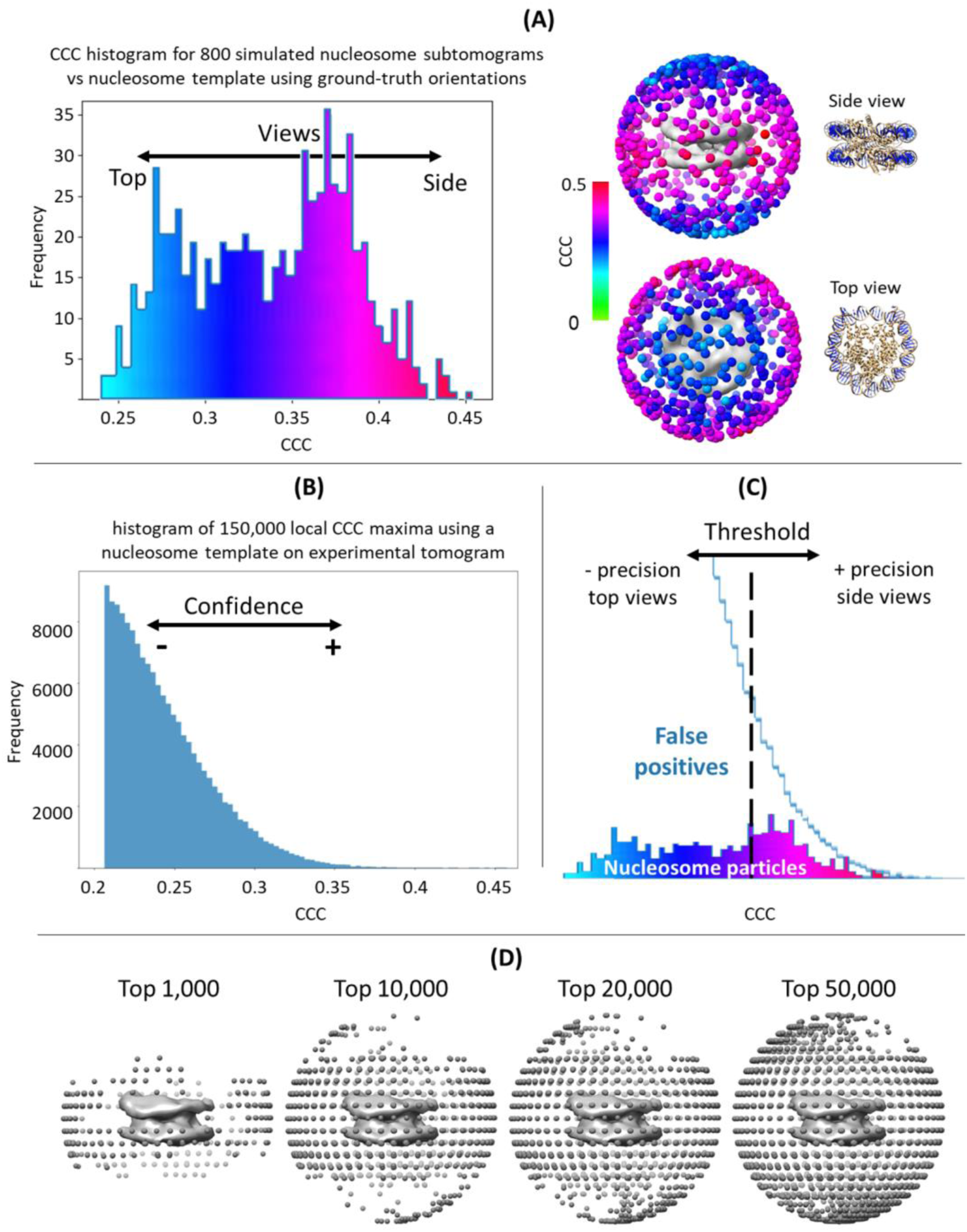
Template matching for nucleosome picking can only extract certain views at an adequate Precision. In this work, we utilized template matching and Template Learning techniques to annotate nucleosomes in cryo-electron tomograms. The angular distribution obtained through averaging of template matching-based annotated particles was found

**Extended Data Tab. 1:** Cryo-ET simulation parameters in Template Learning. The parameters used for domain randomization simulations in this article were based on a combination of the parameters listed in this table.

| Parameter | Values |
| --- | --- |
| Defocus for close-to-focus VPP [microns] | 0, -0.5, -1 |
| Defocus without VPP [microns] | -2.5, -3.25, -4 |
| Electron dose [$e^-/\text{\AA}^2$] | 75, 150 |
| Tilting range [deg] | [-40, 40], [-60, 60] |
| Tilting step [deg] | 2, 4 |
| Ice density [$\text{g}/\text{cm}^3$] | 0.9, 1.1 |

**Extended Data Tab. 2:**
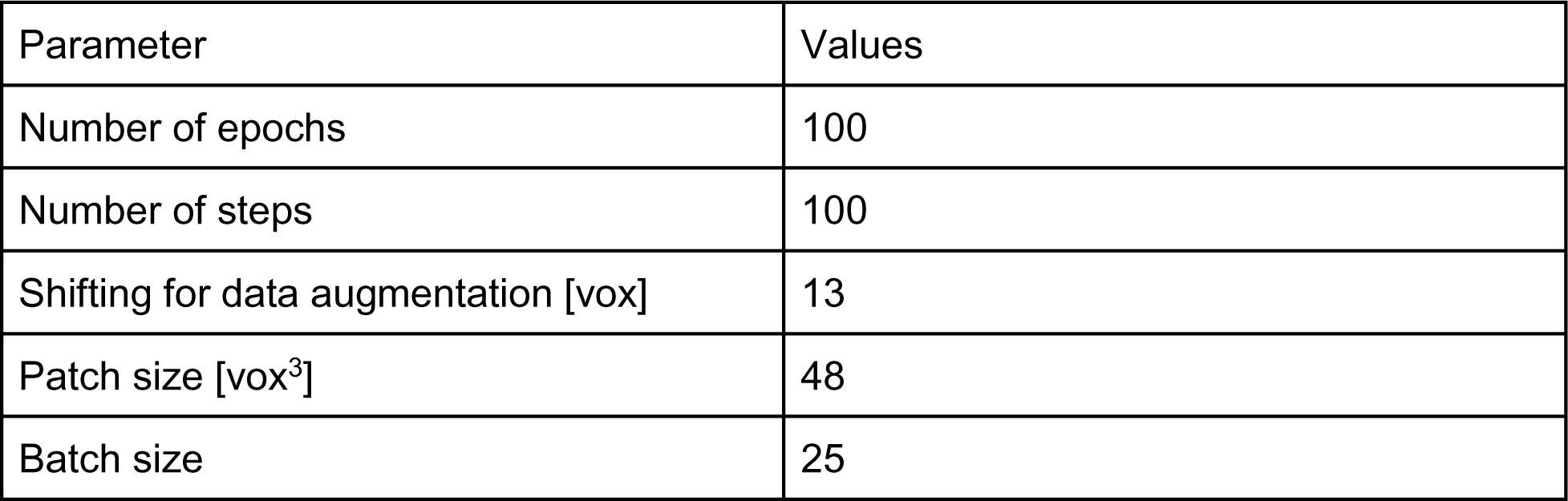
Parameters used for training DeepFinder on Template Learning simulations. Default hyperparameters used to train DeepFinder for the different particle picking experiments.

| Parameter | Values |
| --- | --- |
| Number of epochs | 100 |
| Number of steps | 100 |
| Shifting for data augmentation [vox] | 13 |
| Patch size [$\text{vox}^3$] | 48 |
| Batch size | 25 |

